# Axon Trafficking Counteracts Aberrant Protein Aggregation in Neurons

**DOI:** 10.1101/2025.01.16.633295

**Authors:** Yu Feng, Tongshu Luan, Zhenda Zhang, Wei Wang, Yuanyuan Chu, Sijia Wan, Xiaorong Pan, Jie Li, Yifan Liu, Tong Wang

**Affiliations:** Center for Brain Science, School of Life Science and Technology, Shanghaitech University, Shanghai, 201210, China; Division of Chemistry and Physical Biology, School of Physical Science and Technology, ShanghaiTech University, Shanghai 201210, China

## Abstract

Directed axon trafficking of mRNA via ribonucleoprotein complexes (RNPs) is essential for the proper function and survival of neurons. However, the mechanisms governing RNP transport in axons remain poorly understood. Here, we identify Annexin A7 (ANXA7) as a critical adaptor facilitating the retrograde transport of T-cell intracellular antigen 1 (TIA1)-containing RNPs by linking them to the cytoplasmic dynein. Persistent axonal Ca²⁺ elevation disrupts ANXA7’s linker role, causing the detachment of TIA1 granules from dynein, consequently impairing transport and triggering pathological TIA1 aggregation within axons. Similarly, ANXA7 knockdown decouples TIA1 granules from dynein, severely obstructing trafficking and causing pathological aggregation of TIA1 in axons, which culminates in axonopathy and neurodegeneration both *in vitro* and *in vivo*. Conversely, ANXA7 overexpression enhances trafficking and counteracts aberrant aggregation of TIA1-containing RNPs in axons. Our findings elucidate a novel mechanism underlying RNP axonal transport, highlighting its significance in the biology and pathology of central neurons.

## Introduction

Neurons are exceptionally long and polarised cells, with the axons of projection neurons extending up to one meter ^1^. The functions and survival of these neurons are critically reliant on the axon trafficking of membranes, proteins, and mRNAs to support the demands of remote axonal compartments ^2-5^. mRNAs are packaged with RNA-binding proteins (RBPs) into membrane-less granules called ribonucleoprotein complexes (RNPs), which serve as the actual transport units for axon trafficking ^4,6^. RNPs are transported by molecular motors, primarily kinesin and dynein, either directly via adaptor proteins ^7-10^ or indirectly by tethering to membrane organelles ^11-14^, enabling the rapid, directed and long-range axonal transport along microtubule tracks ^2,6,15^. This motor-driven trafficking is crucial for ensuring the correct localisation and functionality of RNPs within neurons ^4,6,16^.

Accumulating evidence reveals that perturbations in axon trafficking can adversely affect the correct intracellular localisation of RNPs within neurons ^6,17^, leading to the toxic accumulation of RBPs ^16,18^. Specifically, some RBPs are capable of self-assembly into irreversible condensates that potentially play a key pathogenic role ^16,18^. These abnormal aggregates of certain RBPs are considered causal factors for axonopathy and neurodegeneration in diseases such as frontotemporal dementia (FTD) and amyotrophic lateral sclerosis (ALS) ^19-21^. Therefore, understanding how the directed trafficking machinery drives the correct localisation of RNPs in axons, and whether such mechanisms influence RBP aggregation, is crucial for elucidating fundamental axonal biology in healthy and diseased neurons.

One prominent RBP involved in forming pathogenical aggregation is TIA1, which possesses RNA recognition motifs (RRMs) at N-terminus and a prion-like domain (PrLD) at its C-terminus that mediates its prion-like self-aggregation ^22^. In response to axonal injury, TIA1 rapidly forms large RNPs near lesion sites via liquid-liquid phase separation (LLPS), repressing the translation of certain mRNAs and thus suppressing the axon regeneration in a PrLD-dependent manner ^23,24^. Mutations in the PrLD of TIA1 can cause the formation of insoluble (pathological) aggregates and are associated with several neurodegenerative diseases, including FTD and ALS ^25^, and Welander distal myopathy (WDM) ^26^. Furthermore, interaction of wild-type TIA1 with physiological levels of tau/mRNA can lead to toxic tau aggregate formation in neurons ^27^. Reducing TIA1 levels *in vivo* counteracts the tauopathy, protects against neurodegeneration, and prolongs survival of P301S Tau mice, a model for Alzheimer’s disease ^28^. Despite these findings, mechanisms controlling the dynamics of TIA1-containing RNPs in axons remain elusive.

To address this gap, we monitored the movement of TIA1 granules in uni-directional axons of live neurons cultured in microfluidic devices. Our observations revealed that TIA1 granules predominantly move in the retrograde direction. Using mass spectrometry to analyse TIA1 interactors enriched from rat brain lysates, we identified that TIA1 is recruited to the intermediate chain of cytoplasmic dynein, which is motor driving retrograde axon trafficking, via the adaptor protein ANXA7. *In vitro* and axonal trafficking assays demonstrated that elevated Ca^2+^ disrupts ANXA7’s linker function in recruiting TIA1 granules to dynein. ANXA7 knockdown similarly decoupled TIA1 from dynein, impairing its trafficking and leading to the formation of large pathological TIA1 aggregates in live axons. Conversely, ANXA7 overexpression enhanced trafficking efficiency, reducing TIA1 granule aggregates in axons. Furthermore, ANXA7 knockdown in mouse cortex resulted in axonopathy and neurodegeneration, impairing the motor abilities of the mice. Our study uncovers a novel mechanism regulating the dynamic trafficking and aggregation of TIA1-containing RNPs in axons, offering potential avenues for targeting pathogenic protein aggregation underlying neurodegenerative diseases.

## Results

### TIA1 forms RNPs that undergo retrograde trafficking in axons

Known as one of the core proteins composing stress granules (SGs), TIA1 protein suppresses the mRNA translation in many types of cells, incuding the developing neural stem cells ^29-33^, mediated by its PrLD, the LLPS of TIA1 controls the local translation of axonal mRNAs, and thus inhibits the regeneration capacity of injured axons ^24,34^. Besides, LLPS-mediated aggregation of TIA1 also controls tauopathy, thus promoting the degeneration of axon ^27,28^. Given these crutial roles of TIA1 in axon, whether its dynamics is regulated in axons remains largely unknown.

To address this question, in axons of rat hippocampal neurons cultured for 8 days (DIV8), we examined the distribution of endogenous TIA1 and G3BP, which labels stress granules, or SQSTM1/p62, which labels ubiquitinated protein aggregates, and found that the subcellular localisations of these molecules are different (sFig. 1A). TIA1 overlaps with the latter two aggregation markers only in the expanded and beading regions of axons, but not along the unexpanded axon shafts (sFig. 1B). These observations were further supported by results from lattice SIM, revealing the limited co-localisation between TIA1 and either G3BP or p62 in unexpanded axon shafts (sFig. 1C-E). These data agree with previous reports ^24,34^, suggesting that TIA1 granules may have unknown functions in axon shafts.

To explore the potential TIA1 functions in axons, in DIV8 rat hippocampal neurons, we co-expressed TIA1-GFP and CY5-UTP, which label total RNA ^12^, to mark the RNPs formed by TIA1 and RNA. We found that in axons of living neurons, CY5-UTP RNPs are highly mobile with retrograde and anterograde directions (Fig. 1A; sMov. 1). Interestingly, TIA1-GFP preferably overlapping with the RNPs in retrograde direction, reflected by significant higher ratio of TIA1-positive RNPs in the retrograde direction (Fig. 1B). Next, we employed the microfluidic device to separate the axons of cultured hippocampal neurons from the somatodendritic part, as previously described ^35,36^. Then in the uni-directional axon bundles formed within the axon channels, directional trafficking of the flouresent tagged TIA1 granules could be traced and analysed (Fig. 1C). We observed that a substantial number of TIA1-mCherry granules undergo retrograde transport along axons (Fig. 1D; sMov. 2). When categorising the trajectories of these granules into stationary (-0.05 μm/s < speed < 0.05 μm/s), retrograde (speed ≤ -0.05 μm/s), and anterograde (speed ≥ 0.05 μm/s) (Fig. 1E, F), we found that 34.03 ± 2.08% of trajectories were retrograde, significantly higher than the 21.26 ± 1.89% of trajectories that were anterograde (Fig. 1G). These data suggest that most mobile TIA1 granules is retrogradely transported within the axons of living neurons.

**Figure 1.**
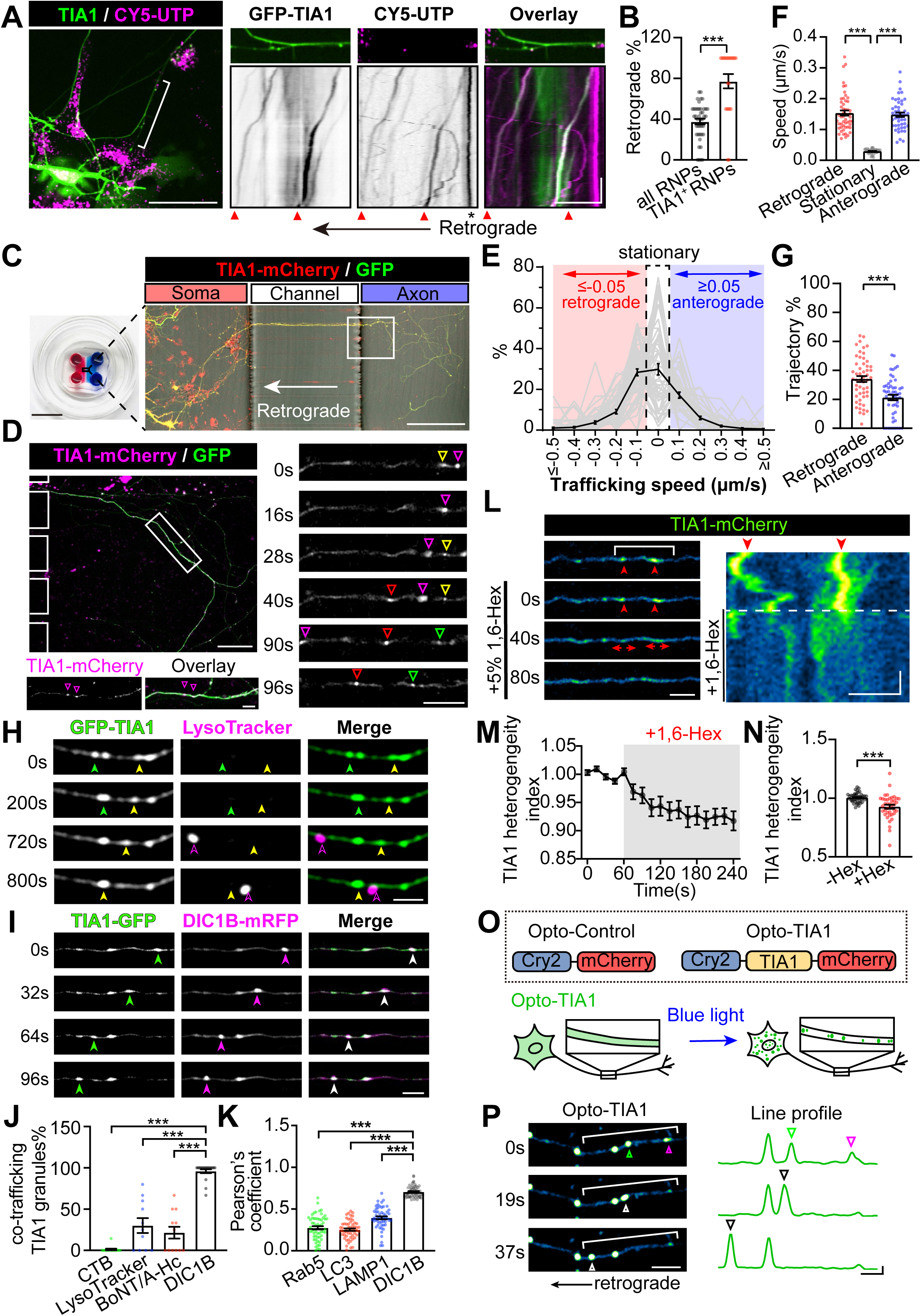
TIA1 forms membrane-less RNPs that undergo retrograde trafficking in axons. **(A)** Live-cell imaging of DIV8 rat hippocampal neurons reveals directed axonal trafficking of RNPs. Kymographs of bracketed axons show retrograde (red triangles) and stationary (black asterisks) RNPs. Scale bar = 50 µm (left), 20 µm (right); y-axis = 180 s. **(B)** Ratio of TIA1-positive RNPs and all RNPs undergoing retrograde trafficking (n = 22, 42). **(C)** Schematic of the microfluidic device. Scale bar = 1 cm (left), 200 µm (right). **(D)** Dynamics of TIA1-mCherry granules in the axons of DIV8 neurons. Magnified boxed regions on the right show distinct TIA1 granules, marked by colored arrowheads. Scale bars = 50 μm and 10 μm. **(E)** Distribution profile of average trafficking speeds for TIA1 granules: retrograde (< -0.05 μm/s), anterograde (> 0.05 μm/s), stationary (-0.05 to 0.05 μm/s) (n = 53). **(F)** Quantification of trafficking speeds of TIA1 granules (n = 53). **(G)** Ratio of retrograde and anterograde TIA1 granules (n = 53). **(H-I)** Time-lapse imaging showing co-trafficking of TIA1 granules with LysoTracker **(H)** and DIC1B-mRFP **(I)** in axons. Scale bar = 5 µm. **(J)** Ratio of TIA1 granules co-trafficking with CTB, BoNT/A-Hc, LysoTracker, and DIC1B-mRFP in axons (n = 14, 12, 11, 18). **(K)** Pearson’s coefficient of endogenous TIA1 granules with indicated markers (n = 56, 56, 53, 56). **(L)** Time-lapse images show the disassembly of axonal TIA1 granules, indicated by red arrows. Right : Kymograph of the indicated axonal segment. Scale bar = 5 µm; y-axis = 10 s. **(M-N)** Quantification shows TIA1 intensity heterogeneity along axons **(M)** before and 180 s post-1,6-Hex **(N)** (n = 40). **(O)** Design of Opto-Control and Opto-TIA1 constructs (top). Schematics illustrating light-induced formation of Opto-TIA1 granules in neurons (bottom). **(P)** Time-lapse images showing formation of Opto-TIA1 granules in axons. Intensity profiles of the highlighted area shown on the right. Moving and fusing granules are indicated by arrowheads. Scale bar = 10 µm (left), 5µm (right); y-axis = 25% (Normalized to (*F_Max_-F_0_*)). Data represent mean ± SEM; two-tailed unpaired *t*-test in **(B, G, N)**; one-way ANOVA in **(F, J, K)**; \*\*\**p*<0.001.

The most well-known mechanism for RNP axon trafficking is the via indirect tethering to membrane organelles proteins ^11-14^. We next explored whether TIA1 granules co-transport with retrograde membranous carriers in axons, using a pulse-chase labeling assay (sFig. 1F), which labels various membraneous organelles originated from axon terminal ^36^. Fluorescently tagged CTB labels signaling endosomes ^35^ (sFig. 1G; sMov. 3), Lysotracker labels lysosomes ^37^ (Fig. 1H; sMov. 3), and the heavy chain of Botulinum neurotoxin (BoNT/A-Hc) labels autophagosomes and multivesicular bodies (MVBs) derived from synaptic vesicles ^38^ (sFig. 1H; sMov. 3). We found that TIA1 granules show limited co-trafficking with signaling endosomes and synaptic vesicle-sourced autophagosomes (sFig. 1G, H) but have a relatively higher overlap with retrograde lysosomes (Fig. 1H, J). Notably, the retrograde motor cytoplasmic dynein, labelled by its neuron-specific intermediate light chain (DIC1B-mRFP) ^39^, demonstrated the highest co-trafficking rates with TIA1 granules in live axons (Fig. 1I, J; sMov. 4). Consistently, in fixed axons of neurons, endogenous TIA1 and these membrane axon carriers also showed limited co-localisation, with the highest overlap observed with dynein (Fig. 1K; sFig. I-K). These results suggest that the axon trafficking of TIA1 granules is unlikely to depend on membranous axon carriers, instead occurring primarily via a direct link with the retrograde motor dynein.

Next, we examined whether the axonal TIA1 granules are membrane-less, by employing two different approaches. First, we used the 5% 1,6-Hexanediol (1,6-Hex) to treat DIV12 neurons, with the dynamics of the TIA1-mCherry monitored under the live-imaging microscopy. We found that the addition of 1,6-Hex causes significant disassembly of TIA1-mCherry granules (Fig. 1L), reflected by the significantly reduced heterogeneity index (Fig. 1M, N), which is defined in sFig. 1L reflects the extent of uneven distribution of fluorescent signals along the thin axons. Second, we used the optoDroplet system ^40^ to construct the light-induced Opto-TIA1, which forms droplets when exposed to blue laser (Fig. 1O). The light-induced Opto-TIA1 droplet were formed within 120 seconds (sFig. 1M; sMov. 5), leading to significantly raised heterogeneity index of Opto-TIA1 (sFig. 1N). We noticed that these Opto-TIA1 granules undergo retrograde trafficking, reflected by the rapid fusion behaviour observed (Fig. 1P; sMov. 6). Results from these two approaches demonstrate that axonal TIA1 granules are indeed membrane-less droplets formed via LLPS.

Collectively, most TIA1 granules in axons are retrogradely transported RNPs that are directly linked to dynein. These TIA1-containing RNPs may have novel functions in axons, distinct from their known SG-related roles.

### ANXA7 is the adaptor linking TIA1 to dynein

To elucidate the mechanisms underlying the direct dynein-driven retrograde trafficking of TIA1 granules in axons, we first compared known TIA1 interactors with those of cytoplasmic dynein intermediate chain 1B (DIC1B), the primary homolog of the dynein intermediate chain 1 in neurons, using data from the BioGrid database. This comparison revealed four shared proteins: ANXA7, CUL3, C2orf44, SMN1 (Fig. 2A; sTab. 1). Next, we expressed recombinant TIA1 in *E. coli* and analysed its interactome using purified GST-TIA1 from P14 rat brain lysates. Following tryptic digestion of GST bead-bound proteins, the resulting peptides were subjected to liquid chromatography-tandem mass spectrometry (LC– MS/MS). Using specific screening criteria (see Methods), we identified 63 proteins enriched in GST-TIA1 compared to GST alone (Fig. 2B; sTab. 2). Gene Ontology (GO) and Kyoto Encyclopedia of Genes and Genomes (KEGG) analysis revealed that these TIA1-associated proteins were enriched in processes related to cytoplasmic stress granule assembly, translation, RNA processing and RNA binding and are associated with neurodegenerative diseases (sFig. 2A; sTab. 3), consistent with TIA1’s established functions ^22,29,31,41-43^. Among the TIA1 interactors, CUL3 and ANXA7 were detected (Fig. 2B). Given ANXA7’s roles in facilitating intracellular trafficking of various organelles ^44-47^ and its confirmed presence in GST-TIA1 pull-downs from P14 rat brain lysates (Fig. 2B, C), we focused on examining whether ANXA7 acts as the linker between TIA1 and dynein.

**Figure 2.**
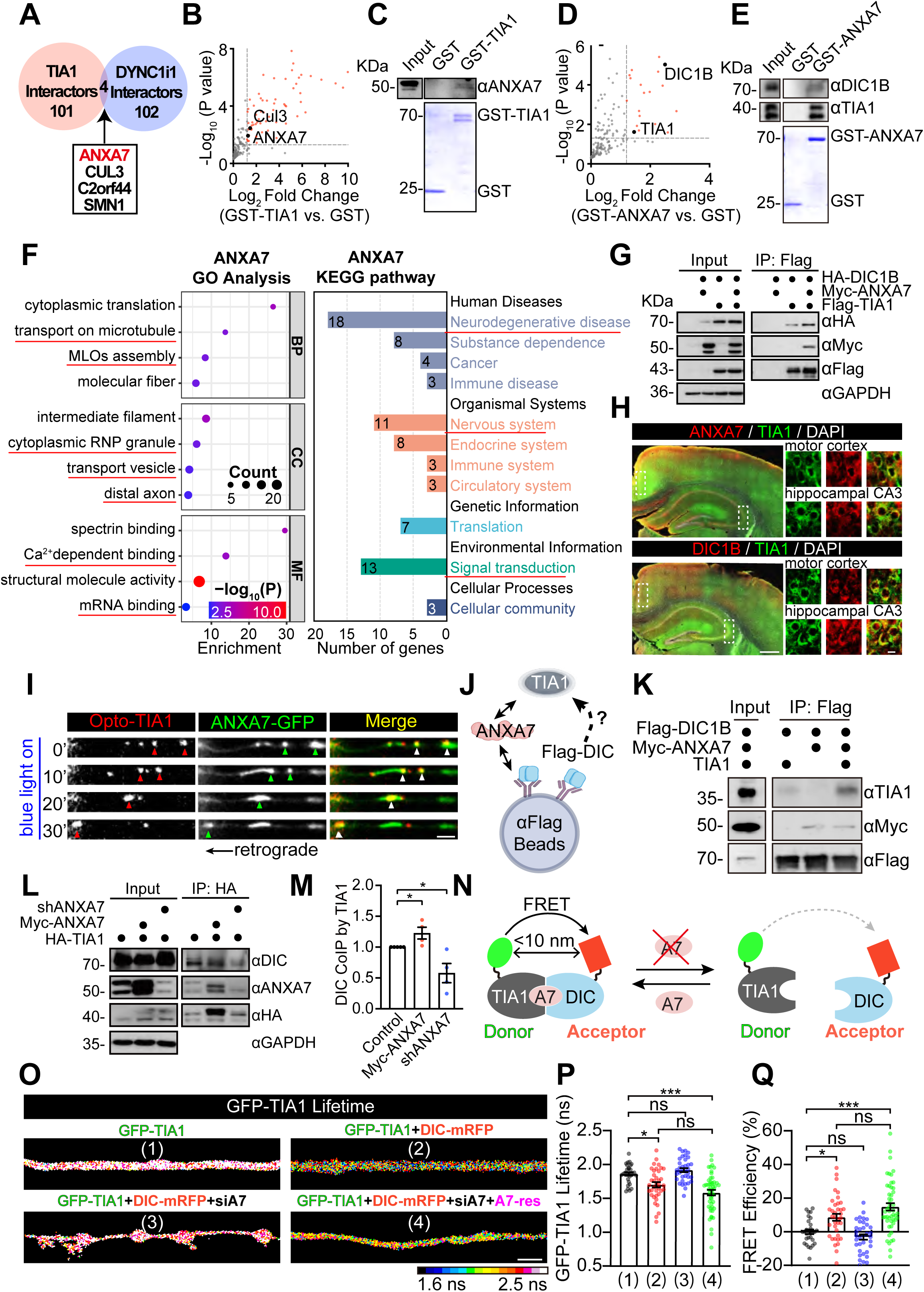
ANXA7 is the adaptor facilitating the interaction between TIA1 and dynein in neurons. **(A)** Venn diagram showing four shared interactors between TIA1 and DIC1B interactomes (DYNC1i1 in BioGrid). **(B, D)** Mass spectrometry analysis of proteins interacting with GST-TIA1 **(B)** or GST-ANXA7 **(D)** in rat brain lysates, using GST tag as a control. Red dots indicate significantly enhanced interactors (*p* < 0.05 and Log2 Fold Change > 1.2). Data from three replicates. **(C, E)** Immunoblots of ANXA7 in proteins pulled down by GST-TIA1 **(C)**; TIA1 and DIC1B in proteins pulled down by GST-ANXA7 **(E)** from rat brain. **(F)** GO and KEGG pathway analysis of GST-ANXA7 interactors, including Biological Processes (BP), Cellular Components (CC), and Molecular Functions (MF). **(G)** Co-IP showing Myc-ANXA7 and HA-DIC1B interact with Flag-TIA1 in HEK293T cells. **(H)** Confocal images of endogenous TIA1 (green) and ANXA7 or DIC1B (red) in the cortex and hippocampus of P34 mouse brain. Scale bars = 500 μm and 10 μm. **(I)** Time-lapse images showing retrograde co-trafficking of light-induced Opto-TIA1 (red) and ANXA7-GFP (green) granules in DIV9 rat hippocampal neurons. Scale bar = 2 µm. Arrowheads indicate co-trafficking. **(J)** *In vitro* protein pull-down assay schematic. **(K)** Purified Myc-ANXA7 protein enhances TIA1 and Flag-DIC interaction, shown by increased TIA1 co-IP’d with Flag-DIC. **(L)** Co-IP assay examining the interaction between endogenous DIC1B and HA-tagged TIA1 using anti-HA magnetic beads in cultured DIV11 rat cortical neurons. The interaction is studied under endogenous ANXA7 knockdown (shANXA7) or Myc-ANXA7 overexpression conditions. **(M)** Quantifying TIA1-DIC1B interaction from **(L)** shows the effects of different ANXA7 levels (n = 5, 4, 4). **(N)** Schematic diagram of FLIM-FRET to examine the affinity between GFP-TIA1 (donor) and DIC1B-mRFP (acceptor) under varying levels of ANXA7. **(O)** Represented images showing colour-coded GFP-TIA1 lifetime in axon shafts of transfected neurons, with lifetime **(P)** and FRET efficiency **(Q)** quantified and compared across indicated groups. Scale bars =2 μm (n = 29, 37, 35, 49). Data represent mean ± SEM; in **(M)** two-tailed unpaired *t*-test; in **(P, Q)** one-way ANOVA. \**p*<0.05, \*\*\**p*<0.001, ns non-significant.

Next, we conducted a pull-down assay using recombinant GST-ANXA7 and P14 rat brain homogenates, with GST-peptide as the negative control. This identified both DIC1B and TIA1 as ANXA7-associated proteins (Fig. 2D; sTab. 2), confirmed by western blot using specific antibodies (Fig. 2E). GO term and KEGG pathway enrichment analyses of the ANXA7 interactome revealed significant enrichment in pathways related to microtubule-dependent transport, membrane-less organelle (MLO) assembly, RNP granules, vesicle trafficking, distal axon function, Ca^2+^-dependent activities, and mRNA binding (Fig. 2F; sTab. 3), suggesting a role for ANXA7 in RNP axonal trafficking. KEGG pathway analysis also highlighted neurodegenerative disease pathways, including ALS, Parkinson’s disease (PD), and Huntington’s disease (HD)—all characterised by early axon degeneration ^21^. These suggest that ANXA7 may function as a crucial linker between TIA1 granules and the dynein complex.

Supporting this hypothesis, Myc-ANXA7 co-immunoprecipitated (co-IP’d) with HA-TIA1 when co-expressed in HEK293T cells (sFig. 2B). Additionally, co-expressed HA-DIC1B co-IP’d with Flag-TIA1, likely facilitated by endogenous ANXA7 in HEK293T cells. Notably, overexpression of Myc-ANXA7 significantly increased the amount of HA-DIC1B co-IP’d with Flag-TIA1 (Fig. 2G), indicating that ANXA7 mediates the interaction between TIA1 granules and the dynein complex.

We explored the expression patterns of endogenous TIA1, ANXA7, and DIC1B in mouse brains using specific antibodies and confocal microscopy. High expression levels of all three proteins were observed in both the cortex and hippocampus, with significant co-localisation in neuronal cell bodies of the motor cortex and hippocampal CA3 region (Fig. 2H). Subcellular distribution in the axons of DIV12 cultured hippocampal neurons was examined using Lattice SIM, revealing significant co-localisation of ANXA7, TIA1, and DIC1B within the axon shafts (sFig. 2C, D). Additionally, light-induced Opto-TIA1 droplets demonstrated co-trafficking in the retrograde direction with the overexpressed ANXA7-GFP granules in axons (Fig. 2I; sMov. 7). Time-lapse Lattice SIM imaging of GFP-TIA1 and ANXA7-mCherry in living neurons showed their co-trafficking along the axon shafts, also highly overlapped in newly formed TIA1/ANXA7 dual positive granules. (sFig. 2E; sMov. 8). These findings suggest that TIA1, ANXA7, and DIC1B likely coexist within the same complex, facilitating the retrograde trafficking of TIA1 granules in axons.

To elucidate whether ANXA7 alone is sufficient to link TIA1 to dynein, we expressed and purified TIA1, Flag-DIC1B (DIC), and Myc-ANXA7 proteins in *E. coli* (sFig. 2F). Using an *in vitro* pull-down assay (Fig. 2J; see Methods), we found that the addition of Myc-ANXA7 significantly increased the amount of TIA1 co-IP’d with DIC (Fig. 2K), suggesting that ANXA7 directly facilitates TIA1 binding to dynein *in vitro*. To validate this role of ANXA7 in neurons, we modulated endogenous ANXA7 levels in DIV11 rat cortical neurons expressing HA-tagged TIA1, and found reducing ANXA7 levels (shANXA7; sFig. 2G) significantly decreased the DIC-TIA1 interaction, while overexpression of Myc-ANXA7 enhanced the interaction (Fig. 2L, M). Furthermore, using Lattice SIM, we assessed the impact of modulating ANXA7 expression on TIA1 recruitment to dynein in the axons of cultured neurons. We found that over-expression of ANXA7-GFP significantly increased the co-localisation of endogenous TIA1 with DIC1B, whereas knockdown of ANXA7 (shA7 1# and 2#) reduced their co-localisation (sFig. 2H, I). These results demonstrate that ANXA7 is sufficient to facilitate the recruitment of TIA1 to dynein in axons.

To investigate whether ANXA7 regulates TIA1 recruitment to dynein in living neurons, we employed fluorescence lifetime imaging microscopy (FLIM). The close proximity between GFP-TIA1 (donor) and DIC1B-mRFP (acceptor) would lead to fluorescence resonance energy transfer (FRET), thereby reducing the fluorescence lifetime of GFP-TIA1, an indication of their interaction. Manipulating ANXA7 levels should affect the FLIM-FRET efficiency between GFP-TIA1 and DIC1B-mRFP if ANXA7 acts as a linker (Fig. 2N). We conducted experiments by knocking down ANXA7 using shRNA (shA7-1#, shA7-2#) and overexpressing a Myc-ANXA7 mutant (Myc-ANXA7-res) resistant to ANXA7 knockdown (sFig. 2J). In axon of hippocampal neurons co-expressing GFP-TIA1 and DIC1B-mRFP (Fig. 2O, (2)), the fluorescence lifetime of GFP-TIA1 (Fig. 2P) was significantly reduced, and FRET efficiency (Fig. 2Q) increased compared to neurons expressing GFP-TIA1 alone (Fig. 2O, (1)), indicating FRET between GFP-TIA1 and DIC1B-mRFP. Conversely, ANXA7 knockdown (Fig. 2O, (3)) significantly prolonged the fluorescence lifetime of GFP-TIA1 (Fig. 2P) and decreased FRET efficiency (Fig. 2Q), suggesting a weakened TIA1-DIC1B interaction. Remarkably, co-expression of Myc-ANXA7-res with siA7 (Fig. 2O, (4)) restored the GFP-TIA1 lifetime and FRET efficiency to near baseline levels (Fig. 2P, Q), demonstrating that ANXA7 is crucial for maintaining the TIA1-DIC1B interaction. Similarly, FLIM-FRET efficiency changes were observed in soma upon ANXA7 level manipulation (sFig. 2K-M). These findings establish that ANXA7 promotes the interaction between TIA1 and dynein in neuronal soma and axons.

### Ca²⁺ elevation disrupts ANXA7-mediated TIA1 recruitment to dynein

ANXA7 contains N-terminal low-complexity domains (LCDs) that potentially mediate LLPS, as illustrated in Fig. 3A. *In vitro* LLPS assays demonstrated that purified ANXA7 protein forms droplets in a concentration-dependent manner (sFig. 3A, B). Droplet formation was significantly enhanced by the addition of the crowding agent PEG8000 (sFig. 3A, C), aligning with previous findings ^48^. The C-terminal region of ANXA7 harbors Ca²⁺-sensitive Annexin Repeats ^49^, which have been shown to trigger LLPS and promote droplet aggregation near plasma membrane (PM) lesions in cancer cells ^45,49^. We then investigated whether intracellular Ca²⁺ affects ANXA7 function in recruiting TIA1 to dynein.

**Figure 3.**
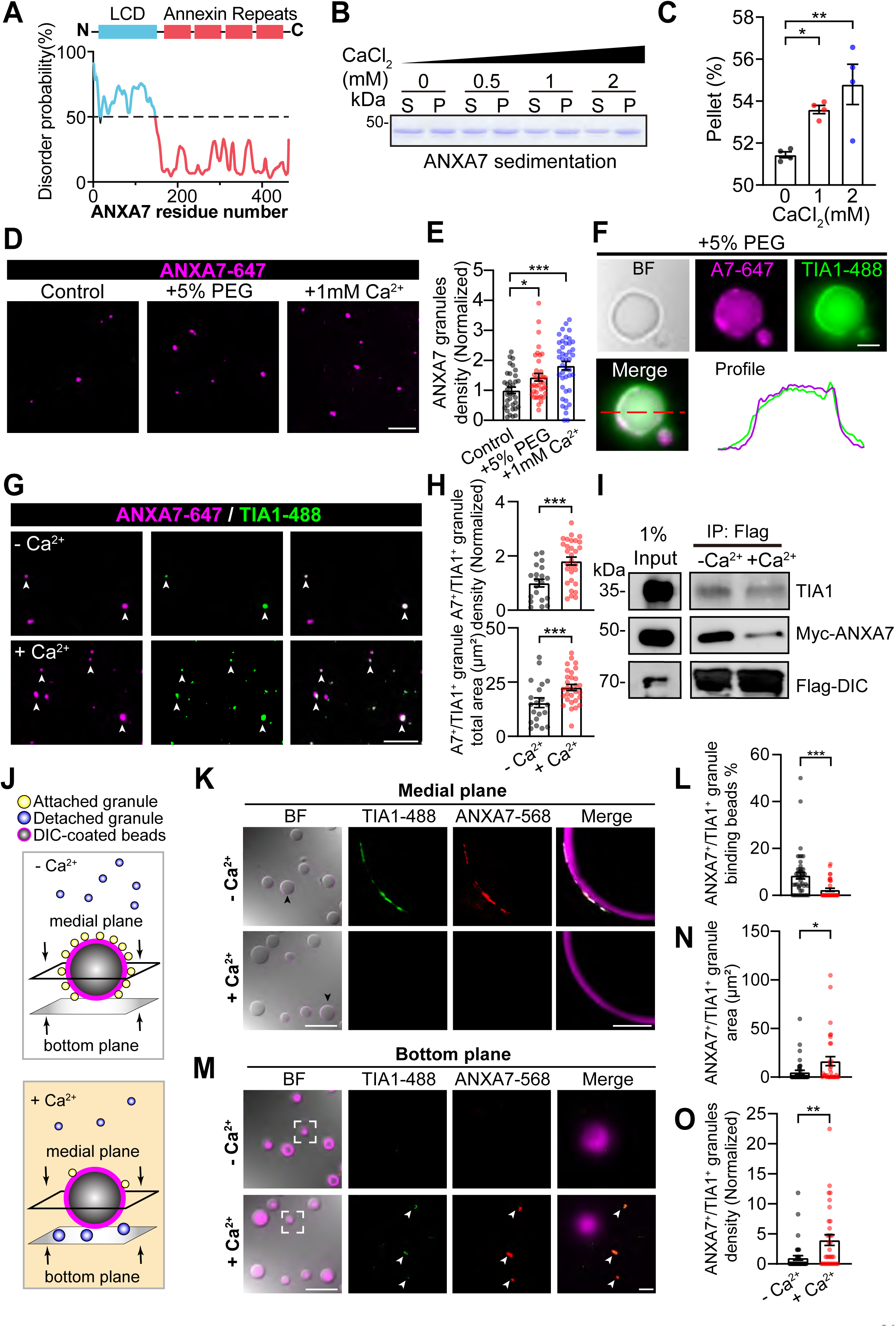
Ca²⁺ promotes ANXA7 aggregation, inhibiting the recruitment of TIA1 to dynein. **(A)** Schematic of ANXA7 domain structure with PrDOS analysis. **(B)** *In vitro* sedimentation assay detected by SDS-PAGE showing the distribution of purified ANXA7 protein (5 μM) between supernatant (S) and pellet (P) at varying Ca²⁺ concentrations. **(C)** Quantification of **(B)** (n = 4). **(D)** *In vitro* phase separation of ANXA7-647 induced by PEG or Ca^2+^. Scale bar = 10 µm. **(E)** Quantification of **(D)** (n = 35, 38, 43). **(F)** *In vitro* droplet formation assay demonstrating co-existence of purified ANXA7-647 and TIA1-488 proteins in droplets induced by 5% PEG. Right: intensity profile along the dashed line in the merged image. Scale bar = 2 µm. **(G)** Confocal microscopy showing increased phase separation of ANXA7-647 and TIA1-488 (both 5 μM) with 1 mM Ca²⁺ addition. Dual-positive droplets indicated by arrowheads. Scale bar = 10 µm. **(H)** Quantification of **(G)** (n = 20, 30). **(I)** *In vitro* interaction assay showing the effect of 1 mM Ca²⁺ on the interaction between purified TIA1, Myc-ANXA7, and Flag-DIC, with Flag-DIC IP’d using anti-Flag beads. **(J)** Schematic diagram of the confocal microscopy-based LLPS assay. See also the Methods for details. **(K, M)** Representative confocal images of the medial plane **(K)** or the bottom plane **(M)** of the DIC-coated beads with or without 1 mM Ca^2+^, showing the ANXA7^+^/TIA1^+^ granules attached **(K)** or detached **(M)** to beads. The boxed regions in the bright field (BF) channel were further amplified in the right panels. Scale bar = 200 µm (left), 20 µm (right). **(L)** Quantification of the percentage of ANXA7^+^/TIA1⁺ granule binding beads in the medial plane shown in **(K)** (n = 46, 35). **(N-O)** Quantifications of the total area **(N)** and density **(O)** of ANXA7^+^/TIA1^+^ condensates in the bottom plane as shown in **(M)** (n = 39, 33). Data represent mean ± SEM; in **(H, L, N, O)** two-tailed unpaired *t*-test; in **(C, E)** one-way ANOVA. \**p*<0.05, \*\**p*<0.01, \*\*\**p*<0.001.

First, we assessed whether Ca²⁺ influences the LLPS of ANXA7 by measuring the sedimentation of purified ANXA7 proteins at various Ca²⁺ concentrations, and found elevated Ca²⁺ levels significantly promoted the segregation of ANXA7 into the pellet fraction (Fig. 3B, C), indicating that Ca²⁺ elevation triggers ANXA7 LLPS, with effects comparable to the addition of 5% PEG *in vitro* (Fig. 3D, E). In the presence of purified TIA1, ANXA7 droplets induced by either 5% PEG or 1 mM Ca²⁺ extensively overlapped with TIA1 droplets (Fig. 3F, G). Notably, TIA1 alone formed droplets *in vitro* with PEG (sFig. 3D-G), but not in response to Ca²⁺ in the absence of ANXA7 (sFig. 3F, H-J), suggesting that the Ca²⁺-triggered formation of TIA1 and ANXA7 droplets (A7⁺/TIA1⁺ droplets) is dependent on ANXA7 (Fig. 3H). These findings indicate that Ca²⁺ elevation induces ANXA7 LLPS, subsequently facilitating TIA1 condensation into the same droplets.

Next, we evaluated the impact of Ca²⁺ elevation on ANXA7’s ability to recruit TIA1 to dynein. Our co-IP assay showed that 1 mM Ca²⁺ significantly reduced the amount of both purified Myc-ANXA7 and TIA1 proteins co-IP’d with Flag-DIC1B (Fig. 3I). To further evaluate the impact of Ca²⁺ on recruitment efficiency, we employed a confocal microscopy-based LLPS assay (see Methods). This allowed us to visualise and quantify A7⁺/TIA1⁺ droplets attached or detached from DIC1B-coated beads across two optical planes: the medial plane, showing the intersection of DIC1B-coated beads with attached A7⁺/TIA1⁺ droplets, and the bottom plane, showing detached condensates (Fig. 3J). We found that 1 mM Ca²⁺ significantly reduced the attachment of A7⁺/TIA1⁺ granules to DIC1B-coated beads in the medial plane (Fig. 3K, L), while significantly increasing the size and number of detached A7⁺/TIA1⁺ droplets in the bottom plane (Fig. 3M-O). These results indicate that Ca²⁺ elevation not only induces the formation of ANXA7⁺/TIA1⁺ droplets but also promotes their detachment from dynein.

### Disruption of ANXA7-mediated trafficking causes TIA1 aggregation in axons

To assess the effect of ANXA7-mediated TIA1 axonal transport under elevated intracellular Ca²⁺ conditions, we used KCl depolarisation to induce significant Ca²⁺ elevation in cultured live neurons^50,51^. We found that application of 56 mM KCl (high K⁺) to DIV11-14 hippocampal neurons expressing the Ca²⁺ sensor GCaMP6f leads to rapidly increase Ca²⁺ levels, represented by the sharp rise in GCaMP6f fluorescence intensity in both somatodendrites (Fig. 4A; sFig. 4A, B; sMov. 9) and axons (Fig. 4A-C), indicating elevated Ca²⁺ levels throughout the neuron. Specifically, the Ca²⁺ intensity in the axon shaft ([Ca²⁺] _axon_) increased by 23 ± 3.4% and remained elevated for over 10 minutes (Fig. 4D). Notably, in expanded axonal regions or “hot spots” (Fig. 4B, arrows), [Ca²⁺]_axon_ levels rose by 4.94 ± 0.5019-fold (Fig. 4E), remaining persistently elevated after depolarisation (Fig. 4C, asterisks). In contrast, 5.6 mM KCl (Low K⁺) did not significantly affect Ca²⁺ levels in either somatodendritic (sFig. 4A, B) or axonal regions (Fig. 4D; sFig. 4C, D).

**Figure 4.**
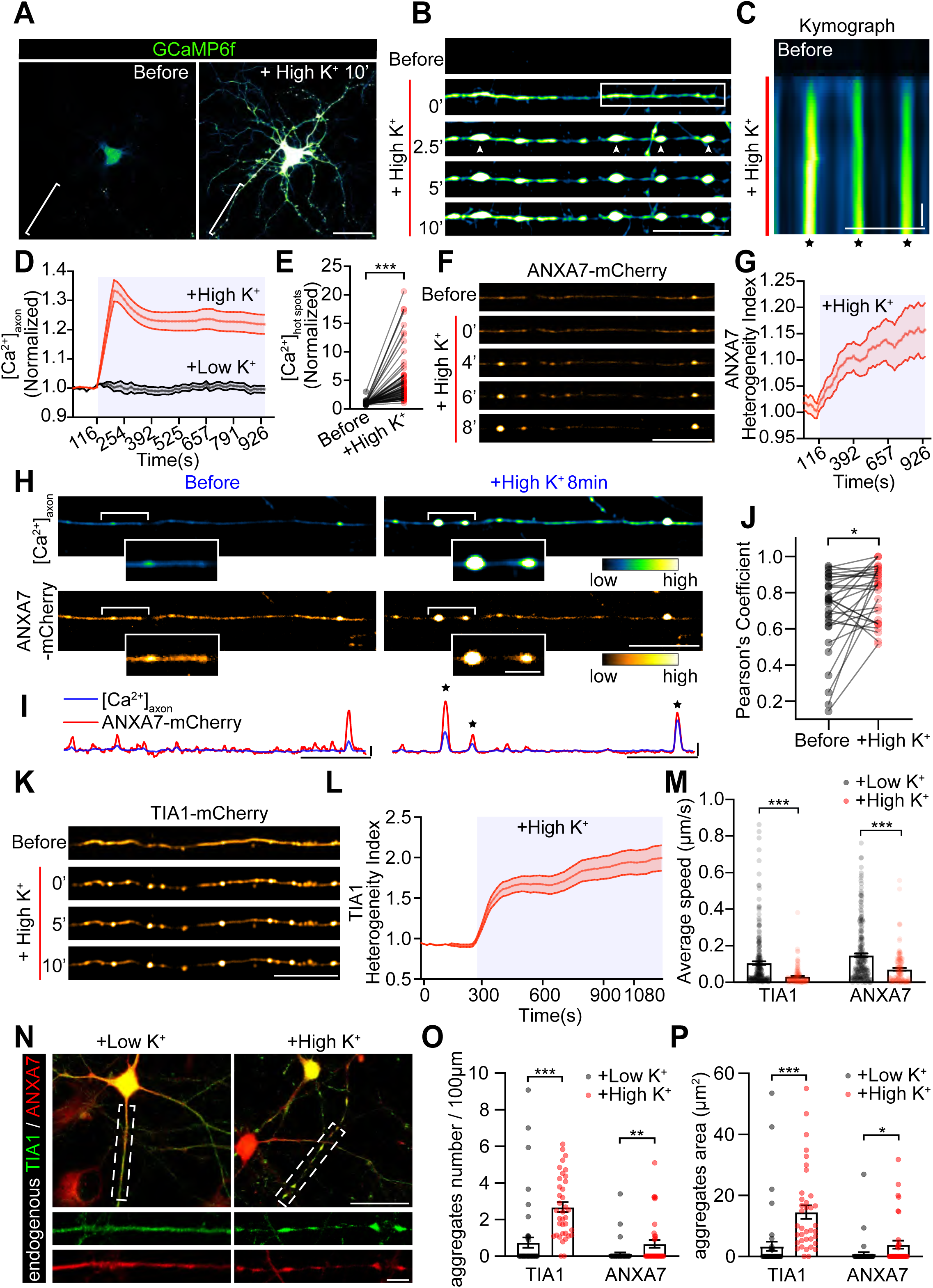
Depolarization-induced Ca²⁺ elevation causes ANXA7 and TIA1 aggregation in axons. **(A)** Live-imaging of DIV13 rat hippocampal neurons expressing GCaMP6f. Scale bar = 50 µm. **(B)** Time-lapse images of axons bracketed in **(A)**, Ca²⁺ elevating "hot spots" indicated with arrows. Scale bar = 20 µm. **(C)** Kymographs of the boxed axon segment in **(B)**, asterisks denote hot spots. Scale Bar = 20 µm; y-axis = 100 s. **(D)** Quantification of [Ca^2+^]_axon_ under different stimulation (Low K⁺: n = 37; High K⁺: n = 72)**. (E)** Quantification of the “hot spots” Ca^2+^ concentration before and after 10 min of high K^+^ stimulation (n = 70). **(F)** Live-imaging of ANXA7-mCherry distribution in axons before and after high K⁺ addition. Scale bar = 20 µm. **(G)** Quantification of ANXA7-mCherry intensity heterogeneity in axons (n = 44). **(H)** Dual-colour images of neurons co-expressing GCaMP6f and ANXA7-mCherry, bracketing axons magnified in insets. Scale bar = 5 µm (left), 20 µm (right). **(I)** Line profiles illustrating fluorescence intensity fluctuations from **(H)**, hot spots denoted by asterisks. Scale Bar = 20 µm; y-axis = 25% (Normalized to (*F_Max_-F_0_*)). **(J)** Pearson’s coefficient showing the correlation between [Ca^2+^]_axon_ and ANXA7-mCherry before and after high K^+^ stimulation (n = 30). **(K)** Live-imaging graphs of TIA1-mCherry distribution in axons before and after high K⁺ addition. Scale bar = 20 µm. **(L)** Quantification of TIA1-mCherry intensity heterogeneity in axons (n = 58). **(M)** Quantification of the average speed of TIA1 and ANXA7 granules under indicated conditions (TIA1: n = 236, 127; ANXA7: n = 241, 134). **(N)** Confocal images of endogenous TIA1 and ANXA7 distribution in axons of DIV12 cultured rat hippocampal neurons after 10 min low or high K⁺ stimulation. Boxed axons are magnified at the bottom. Scale bar = 50 µm (top), 10 µm (bottom). **(O-P)** Quantification of **(N)**, showing the number **(O)** and total area **(P)** of TIA1 or ANXA7 granules per 100 μm axon under the indicated stimulations (n = 48, 37 for both **(O)** and **(P)**). Data represent mean ± SEM; two-tailed unpaired *t*-test **(M, O, P)**; two-tailed paired *t*-test **(E, J)**; \**p*<0.05, \*\**p*<0.01, \*\*\**p*<0.001.

Having established a live-cell model of depolarisation-induced, persistent Ca²⁺ elevation in axonal hot spots, we next examined its effect on ANXA7-mediated axonal trafficking of TIA1. ANXA7-mCherry accumulated in these regions, coinciding with Ca²⁺ elevation (Fig. 4F, G). In neurons co-expressing GCaMP6f and ANXA7-mCherry, co-localisation of Ca²⁺ elevation and ANXA7-mCherry accumulation was observed (Fig. 4H; sMov. 10), confirmed by overlapping peaks in fluorescence line profiles (Fig. 4I, asterisks) and a significant increase in Pearson’s coefficient (Fig. 4J). The aggregation of ANXA7 in these Ca²⁺-elevated axonal hot spots resembled the Ca^2+^-induced ANXA7 aggregates previously seen in cancer cells ^45^. Notably, TIA1-mCherry also significantly accumulated in expanded axonal regions following depolarisation (Fig. 4K, L). These data suggest that Ca²⁺ elevation promotes the formation of ANXA7 and TIA1 aggregates in axonal hot spots with persistent [Ca²⁺]_axon_ elevation.

To further investigate the effect of Ca²⁺ elevation on TIA1 granule trafficking, we measured the movement speed of TIA1-mCherry and ANXA7-mCherry granules, and found that 10 minutes after depolarisation, the trafficking speed of both granules significantly decreased (Fig. 4M). Additionally, 10 minutes of high K⁺ depolarisation led to a significant increase in both the number and size of endogenous TIA1 and ANXA7 granules in DIV11 hippocampal neurons compared to the low K⁺-treated group (Fig. 4N-P). These results indicate that depolarisation-induced persistent Ca²⁺ elevation in axonal "hot spots" promotes ANXA7 aggregation, impedes ANXA7-mediated TIA1 trafficking, and triggers TIA1 aggregation in axons.

### ANXA7-mediated TIA1 trafficking is crucial for maintaining axon integrity

To investigate whether ANXA7-mediated TIA1 axon trafficking affects axon health, we conducted gain- and loss-of-function experiments in DIV12 cultured hippocampal neurons. Our results showed that manipulating ANXA7 levels significantly altered the extent of TIA1 granules aggregation within axons (Fig. 5A). Specifically, overexpression of ANXA7 led to a reduction in the formation of large TIA1 granules (≥2 μm²) (Fig. 5B, C), whereas downregulation of endogenous ANXA7 using shRNA (shANXA7) resulted in a significant increase in the number of large TIA1 granules within axons (Fig. 5A, arrows; Fig. 5B, C). Overexpression of an ANXA7 variant resistant to shRNA knockdown (shA7+A7-res) reversed the phenotype caused by ANXA7 knockdown, reducing the size of TIA1 granules in the axon (Fig. 5B, C). Consistent results were observed with co-expressed TIA1-mCherry granules, showing a reduced number of large granules in ANXA7-overexpressing neurons, and an increase in large TIA1-mCherry puncta when endogenous ANXA7 was knocked down (sFig. 5A, B). These findings indicate that ANXA7 is critical in preventing TIA1 aggregation within axons.

**Figure 5.**
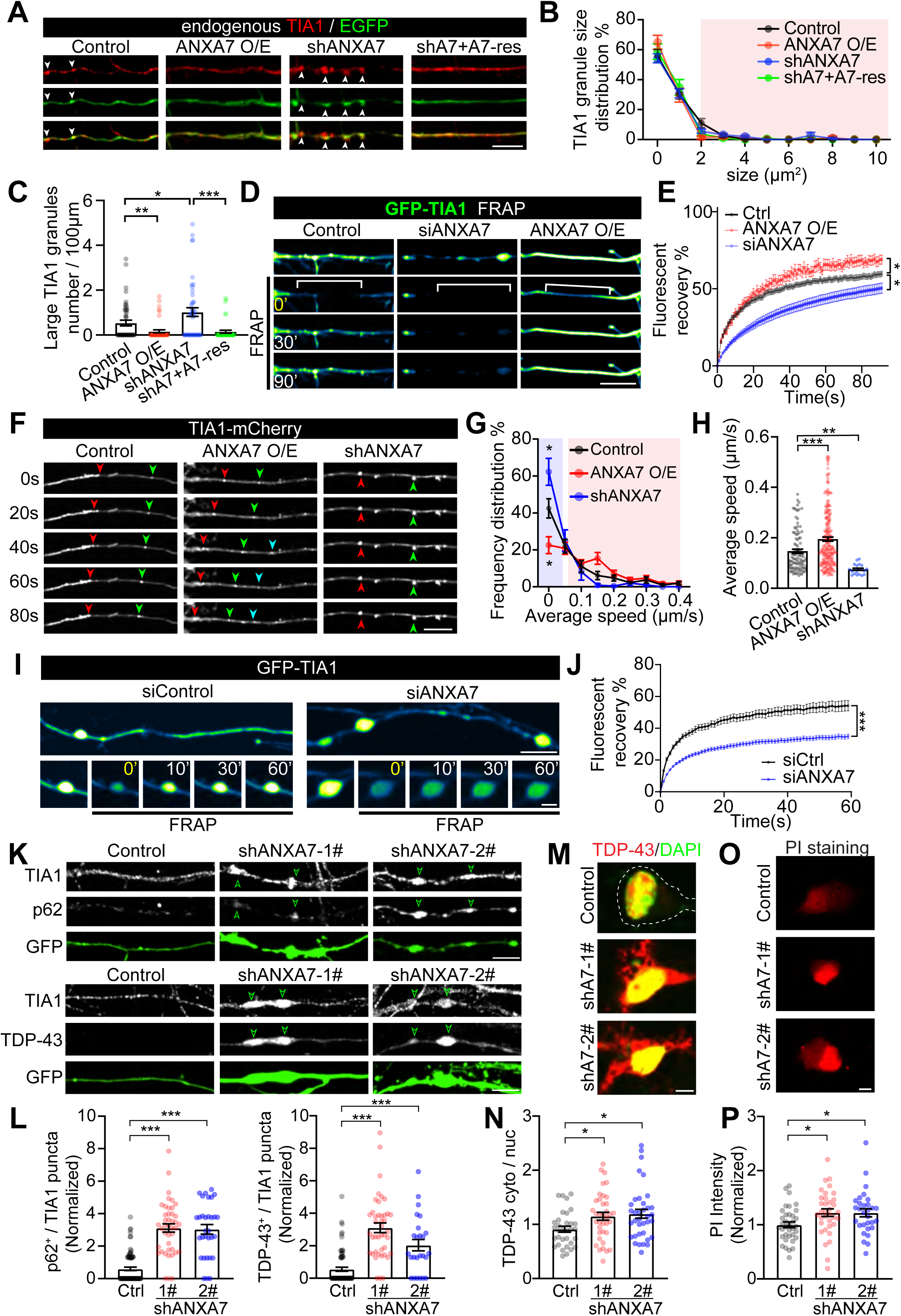
ANXA7 is essential for TIA1 axon trafficking and phase separation in neurons. **(A)** Representative images of TIA1 granules in DIV12 rat hippocampal neuron axons under different ANXA7 expression levels. Arrowheads indicate TIA1 granules. Scale bar = 10 µm. **(B)** Quantification of TIA1 granule size from **(A)**. The red shadow indicates large granules (≥ 2 μm^2^, circularity 0.6-1) (n = 43, 43, 46, 40). **(C)** Ratio of large TIA1 granules (n = 47, 46, 47, 44). **(D)** Time-lapse images of GFP-TIA1 FRAP in axons. Brackets indicate photobleached segments. Scale bars =10 μm. **(E)** Quantification of FRAP curves from **(D)** (n = 90, 27, 69). **(F)** Time-lapse images showing retrograde trafficking of GFP-TIA1 granules. Different coloured arrowheads denote distinct granules. Scale bar = 10 µm. **(G)** Distribution of the GFP-TIA1 granule speeds from **(F)**. Blue shadow indicates stationary granules (≤ 0.05 μm/s), red shading indicates mobile (> 0.05 μm/s) granules (n = 21, 23, 23). **(H)** Average speed of moving GFP-TIA1 granules (speed > 0.05 μm/s) from **(G)** (n = 102, 214, 21). **(I)** Time-lapse images of FRAP for stationary GFP-TIA1 granules in axons. Scale bar = 15 µm (top), 5 µm (bottom). **(J)** Quantification of FRAP curves from **(I)** (n = 160, 199). **(K)** Distribution of endogenous TIA1 with p62 (top) or TDP-43 (bottom) in axons. GFP depicts axon morphology. Scale bar = 10 µm. **(L)** Quantification from **(K)** showing the number of TIA1 puncta co-localized with p62 (left) or TDP-43 (right) per 100 μm axon. (p62^+^: n = 79, 42, 31; TDP43^+^: n = 70, 42, 25). **(M)** Distribution of endogenous TIA1 and TDP-43 in the cytoplasm and nucleus in neurons. Dash line in “Control” depicts soma morphology. Scale bar = 5 µm. **(N)** Quantification of **(M)** (n = 33, 37, 36). **(O)** PI staining images identifying dead cells in the indicated groups. Scale bar = 5 µm. **(P)** Quantification of **(O)** (n = 36, 35, 30). Data represent mean ± SEM; two-tailed unpaired *t*-test in **(C, J)**; one-way ANOVA in **(E, G, H, L, N, P)**; \**p*<0.05, \*\**p*<0.01, \*\*\**p*<0.001.

Next, we determined how ANXA7 prevents the formation of large TIA1 aggregates in axons. Since phase separation capacity is closely related to the molecular mobility of RNA-binding proteins ^52^, we assessed whether ANXA7 modulates the overall mobility of TIA1 molecules in axons by measuring fluorescence recovery after photobleaching (FRAP) of GFP-TIA1 along long axonal segments (∼30 μm) (Fig. 5D; sMov. 11). FRAP rates were enhanced in ANXA7-overexpressing axons but reduced in ANXA7 knockdown axons (Fig. 5E). The FRAP efficiency across long axonal segments reflects the combined effects of trafficking- and diffusion-dependent mobility of TIA1 molecules within the axon. To distinguish the contributions of these two mechanisms, we first evaluated TIA1 granule trafficking by automatically tracking TIA1-mCherry granules and comparing their average speed along axons (Fig. 5F; sMov. 12). ANXA7 knockdown significantly increased the ratio of immobile TIA1 granules (Fig. 5G, blue shaded), whereas ANXA7 overexpression resulted in a higher ratio of mobile granules (Fig. 5G, red shaded). The average speed of mobile TIA1 granules was faster in ANXA7-overexpressing axons and slower in those with reduced ANXA7 (Fig. 5H). Similarly, the trafficking efficiency of light-induced Opto-TIA1 granules was significantly reduced in ANXA7 knockdown neurons (sFig. 5C, D), although their formation capacity remained unaffected (sFig. 5C, E). These results suggest that ANXA7 facilitates active axonal trafficking of TIA1 granules.

We then examined the mobility of TIA1 molecules within the large, immobile granules frequently observed in ANXA7 knockdown neurons, using FRAP assay (Fig. 5I; sMov. 13), and found that GFP-TIA1 mobility within these granules was dramatically reduced compared to control neurons (Fig. 5J). This indicates that these larger, immobile TIA1 granules, induced by ANXA7 downregulation, resemble condensed aggregates. Notably, axons of neurons with ANXA7 knockdown using two distinct shRNA sequences (shANXA7-1# and shANXA7-2#) exhibited a significant increase in TIA1 aggregates (sFig. 5F, G), along with multiple axonal swellings and bead-like morphologies (sFig. 5F, arrowheads; sFig. 5H), indicative of focal axonal swellings (FAS), a hallmark of axonopathy ^53-55^. IF staining revealed that TIA1 aggregates in FAS regions significantly overlap with SQSTM1/p62 and TDP-43, which are markers of pathological aggregations (Fig. 5K, L) ^56-58^. Additionally, ANXA7 downregulation significantly increased the cytoplasmic distribution of TDP-43 from the nucleus (Fig. 5M, N) and resulted in a higher proportion of neurons with unhealthy nuclei (Fig. 5O, P; sFig. 5I, J), suggesting exacerbated neurodegeneration in the absence of ANXA7.

Collectively, these findings from primary neurons demonstrate that the loss of ANXA7-mediated TIA1 axon trafficking promotes the aggregation of large TIA1 granules, leading to axonopathy and neurodegeneration. This underscores the crucial role of ANXA7 in maintaining axon integrity by ensuring the proper recruitment of TIA1 granules to dynein.

### ANXA7 down-regulation induces neurodegeneration in the mouse motor cortex

To further explore the *in vivo* function of ANXA7-mediated mechanism in axon integrity, we down-regulated the ANXA7 expression in neurons of the motor cortex of neonatal mice by intra-cerebroventricular injection (ICV) of two effective shRNA sequences (3# or 4#) (sFig. 6A) delivered by AAV-U6-shANXA7-hSyn-GFP in postnatal day 1 (P1) neonatal mice (Fig. 6A). The AAV-hSyn-GFP was used as control. Then, as illustrated in Fig. 6B, from P56 to P58, mice underwent three rotarod training sessions, followed by a latency-to-fall test on P59, before being sacrificed on P60. We found that ANXA7 knockdown mice exhibited significantly impaired motor capacity, evidenced by their shorter latencies on the rotarod compared to control mice (Fig. 6C). Motor information was delivered from the upper motor neuron (layer V) in the motor cortex to spinal motor neurons via the long-range axon projections in corticospinal tract (CST), which lies in the lateral of spinal cord (Fig. 6D). We observed that ANXA7 knockdown led to a thinner layer V in the motor cortex (M1 and M2 region^59^) (Fig. 6E, F), suggesting loss of upper motor neurons. Morphological analysis of axons in the lateral CST revealed a higher percentage of axons exhibiting typical FAS morphology (Fig. 6G, H), indicative of axon degeneration. We also examined TIA1 distribution in the soma of ANXA7-knockdown neurons in layer V, finding increased cytosolic TIA1 aggregates co-localised with SQSTM1/p62 and TDP-43 (Fig. 6I-L; sFig. 6B, C), suggesting pathological aggregation.

**Figure 6.**
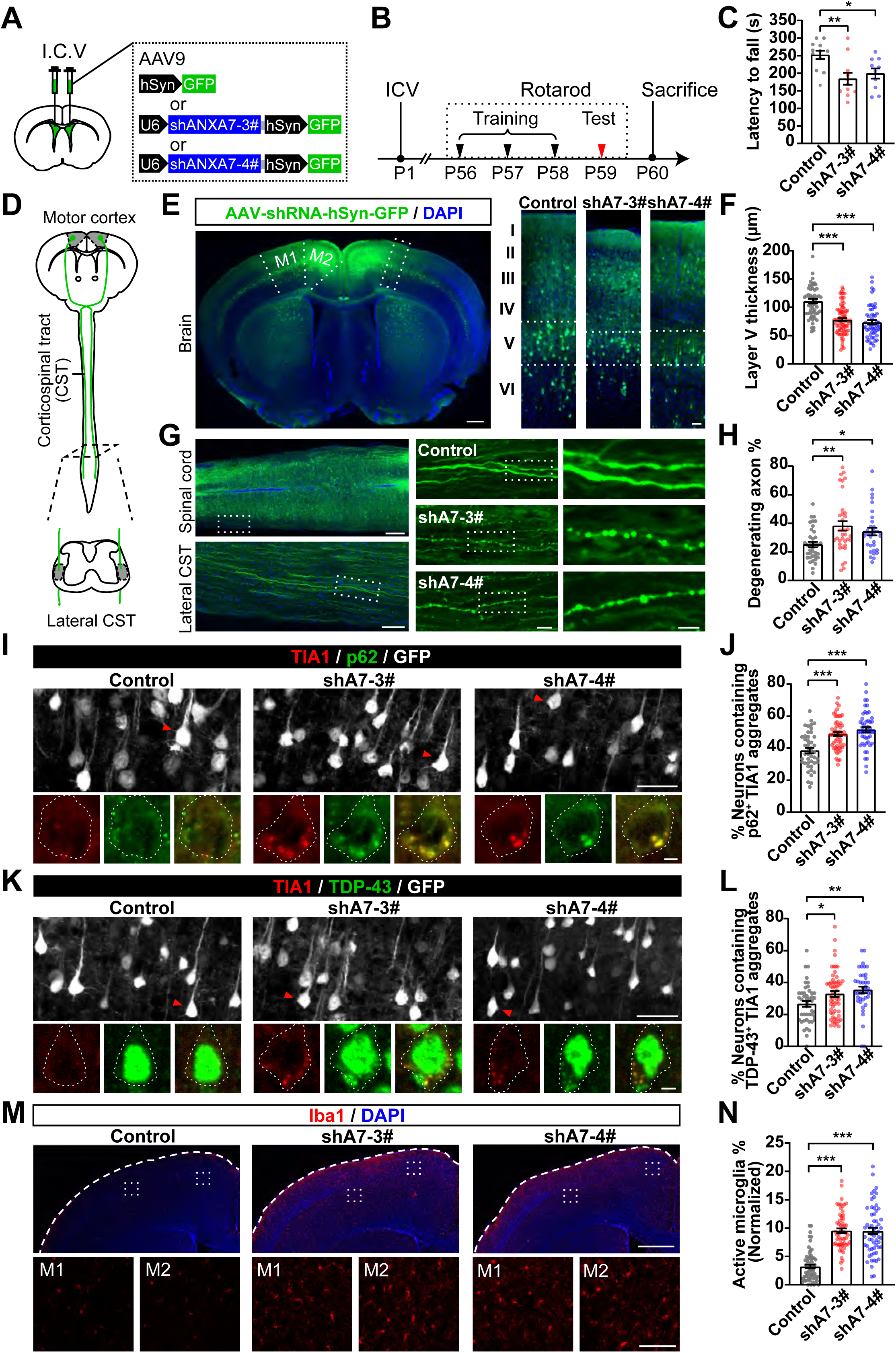
Down-regulation of ANXA7 causes neurodegeneration in mice motor cortex. **(A)** Diagram of intra-cerebroventricular (ICV) injection sites in P1 mouse pups for delivering shANXA7 (shA7-3# and 4#). **(B)** Experimental timeline. P1: ICV of AAVs; P56-P59: Rotarod training and test; P60: sacrifice and tissue IF staining. **(C)** Rotarod probe test results showing latency to fall (n = 12, 11, 10). **(D)** Diagram of upper motor neuron projection pathways, showing somas in cortical layer V and descending axons in the corticospinal tract (CST). **(E)** Confocal images of P60 mouse motor cortex, with the M1 and M2 indicated in the right panel and thickness of layer V marked between dashed lines in the left panels. Scale bar = 500 µm (left), 50 µm (right). **(F)** Quantification of layer V thickness in the cortex (n = 60, 83, 56). **(G)** Confocal images of lateral CST in P60 mouse spinal cord showing descending axons of infected cortical neurons. Magnified boxed regions detail individual axon morphology. Scale bar = 500 µm (left top), 100 µm (left bottom), 20 µm (middle) and 10 µm (right). **(H)** Proportion of degenerating axons with beading morphology (n = 37, 36, 33). **(I, K)** Confocal images of P60 mouse cortex showing TIA1 and p62 **(I)** or TDP-43 **(K)** IF in layer V neurons. Infected neurons marked by GFP expression, the neurons pointed by red arrows are amplified in lower pannels. Dotted lines depict soma shapes. Scale bar = 50 µm (top), 5 µm (bottom). **(J)** Ratio of neurons containing p62^+^ TIA1 aggregates (n = 45, 55, 50). **(L)** Ratio of neurons containing TDP43^+^ TIA1 aggregates (n = 45, 58, 41). **(M)** Confocal images of P60 mouse motor cortex showing the density of active microglia, detected by Iba1 staining. Boxed areas are amplified in the bottom panels. Scale bar = 200 µm (top) and 100 µm (bottom). **(N)** Ratio of Iba1^+^ active microglia (n = 60). Data represent mean ± SEM and from 3 mice; one-way ANOVA in **(C, F, H, J, L, N)**; \**p*<0.05, \*\**p*<0.01, \*\*\**p*<0.001.

Furthermore, we found that in the motor cortex of ANXA7 knockdown mice, the number of reactive microglia, indicated by the density of Iba1^+^ cells, was significantly increased compared to controls (Fig. 6M, N), reflecting the occurrence of neurodegeneration in these regions ^60^. These results demonstrate that ANXA7 knockdown in the mouse motor cortex leads to abnormal TIA1 aggregation in neurons, resulting in axonopathy and neurodegeneration *in vivo*. This further supports the notion that ANXA7-mediated trafficking of TIA1 granules is crucial for maintaining the integrity of long projection neurons in the CNS.

Conclusively, our study uncovered a direct mechanism underlying the axonal transport of RNPs in CNS neurons, highlighting its role in maintaining axonal integrity. As illustrated in Fig. 7, ANXA7 links TIA1 granules to dynein motor, enabling their long-range retrograde axon trafficking. Persistent Ca²⁺ elevation disrupts this linkage, causing detachment of ANXA7/TIA1 granules from dynein, leading to TIA1 aggregation within focal axonal regions. Similarly, ANXA7 knockdown decouples TIA1 granules from dynein, hindering their axon trafficking and promoting their pathological aggregation, which subsequently leads to axonopathy and neurodegeneration both *in vitro* and *in vivo*. Conversely, ANXA7 overexpression enhances axon trafficking and mitigates TIA1 axonal aggregates, suggesting a therapeutic potential for the removal of pathological aggregates and alleviation of neurodegeneration.

**Figure 7.**
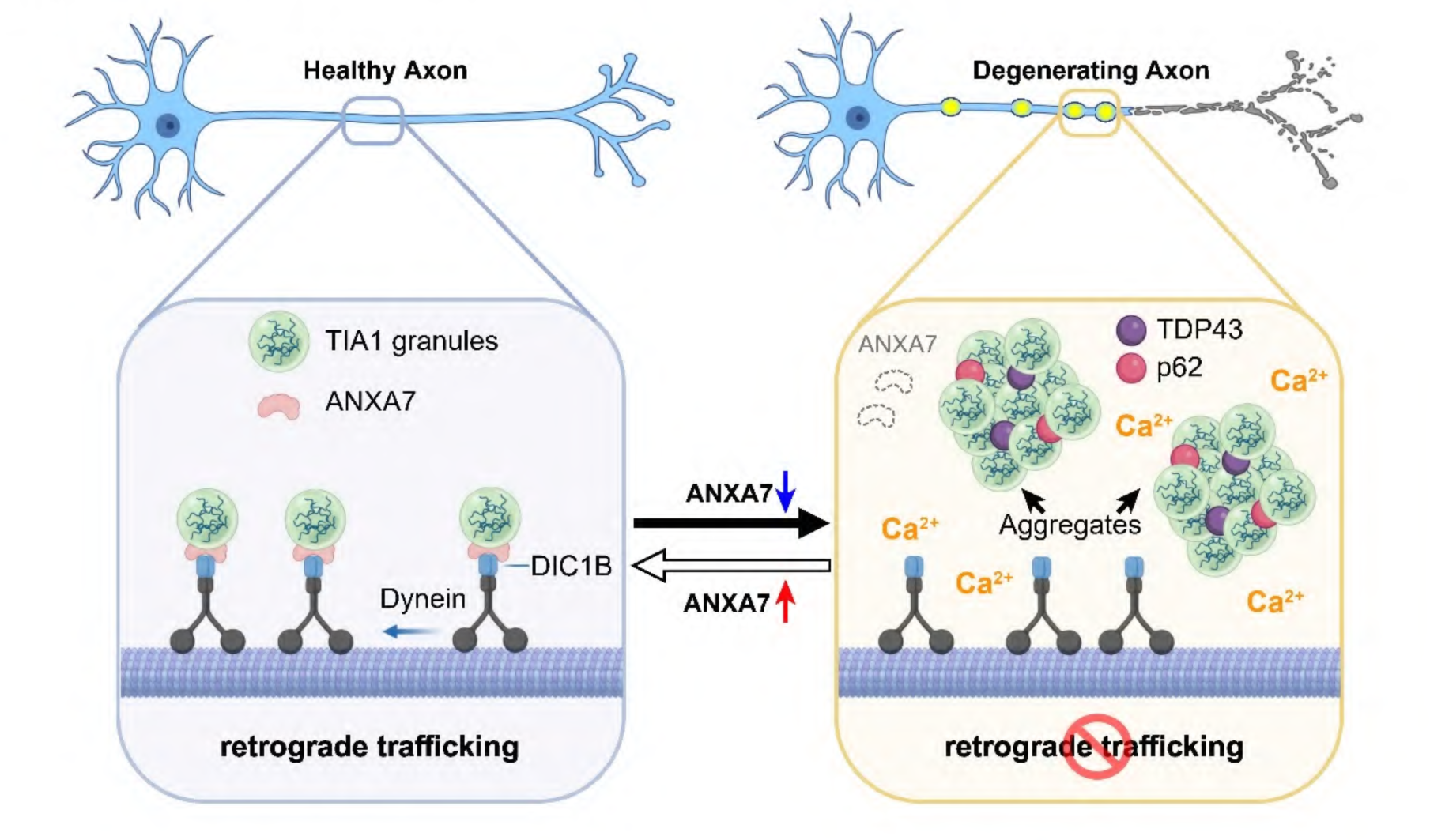
ANXA7-mediated axon trafficking of TIA1 granules counteracts aberrant aggregation in neurons. **Left:** ANXA7 links TIA1 granules to the dynein subunit DIC1B, facilitating their long-range retrograde trafficking in the axon. This process is crucial for maintaining the health of the axon. **Right:** Ca²⁺ elevation disrupts ANXA7’s linker function, causing its detachment from dynein, leading to TIA1 aggregation in focal axonal regions where Ca²⁺ surges persist. ANXA7 knockdown similarly decouples TIA1 granules and dynein, reducing axonal trafficking and resulting in the local accumulation and pathological aggregation of TIA1 granules. These aggregates contribute to axonopathy and neurodegeneration both *in vitro* and *in vivo*. Conversely, ANXA7 overexpression enhances axon trafficking and counteracts aberrant TIA1 aggregate formation.

## Discussion

The directed axon trafficking system delivers RNPs to meet the dynamic demands for proteins and mRNAs in polarised and extended neurons ^6,61^. Some of these RNPs contain RBPs with PrLDs that are prone to forming toxic fibrils, leading to neurodegeneration ^27,28^. Not surprisingly, defects in axon trafficking are closely associated with the abnormal aggregation of these fibril-forming RBPs ^3,17^. Nevertheless, it remains unknown whether the directed transport machinery is involved in preventing RBP aggregation in axons, which is an early indicator and causal factor for neurodegeneration. In this study, we revealed that dynein-driven RNP trafficking counteracts TIA1 aggregation in axons, providing a potential strategy to target pathogenic aggregations underlying neurodegenerative diseases.

### ANXA7 directly links TIA1-containing RNPs to dynein

Two primary mechanisms underlie the bidirectional trafficking of RNPs: the indirect mechanism, in which RNPs are tethered to membranous organelles such as lysosomes ^11^ and endosomes ^12,62^, and the direct mechanism, where the RBP components of the RNPs are directly linked to motors via adaptor proteins, for instance, adenomatous polyposis coli (APC) interacts with kinesin-associated protein 3 (KAP3), an adaptor for the anterograde motor kinesin-2 ^8^; splicing factor proline/glutamine-rich (SFPQ) interacts with kinesin light chain 1 (KLC1) of kinesin family member 5A (KIF5A) ^63^. Although the retrograde motor dynein was identified to drive RBP trafficking via binding to adaptor proteins BicD1 and RBP Egalitarian (Egl) ^9,10^, to date, no direct mechanism for dynein-mediated axonal transport of RNPs has been identified.

In neurons expressing TIA1-GFP and RNA labeled with CY5-UTP, we unexpectedly observed that, unlike the entire pool of CY5-UTP-labeled RNPs, which exhibit bidirectional transport, TIA1-containing RNPs predominantly move in a retrograde direction (Fig. 1A-B). This finding was validated in uni-directional axons of neurons cultured in microfluidic devices (Fig. 1C-G). This retrograde bias suggested a link to dynein, the motor driving retrograde transport from axon tips to the soma ^64^. Through mass spectrometry of TIA1 interactors from mouse brain lysates, we identified ANXA7, a Ca^2+^-regulated protein, that interacts with both TIA1 and the dynein subunit DIC1B. Live-cell confocal microscopy and structured illumination microscopy (SIM) validated significant co-localisation and co-transportation of TIA1, ANXA7, and dynein in primary neurons. Further, co-IP and *in vitro* pull-down experiments demonstrated that ANXA7 strengthens the interaction between TIA1 and DIC1B. Moreover, FLIM-FRET experiments in live neurons showed ANXA7 expression controls their interaction. Our data suggest ANXA7 mediates the direct interaction between TIA1 and dynein, facilitating the retrograde transport of TIA1 granules in axons. This discovery reveals a direct dynein-driven RNP transport mechanism in neuronal axons, filling the gap in understanding how RNPs are linked to the retrograde trafficking machinery—a process that has vital functional implications in the nervous system ^6,65-67^.

### Ca^2+^ overload inhibits the ANXA7-mediated recruitment of TIA1 to dynein

With an N-terminal proline-rich LCD domain, the ANXA7 undergoes LLPS, which is triggered by Ca^2+^ elevation in cancer cells ^48^. Meanwhile ANXA7 is a Ca^2+^-sensitive protein with C-terminal Annexin repeats, which mediates its Ca^2+^-triggered phospholipid binding to the plasma membrane (PM) ^45,48,49^. Highly expressed in neurons, ANXA7 is known as a positive regulator of synaptic vesicle release and postsynaptic N-methyl-d-aspartate (NMDA) receptor trafficking ^68^. But whether AXNA7 plays any role in axon trafficking, and whether such function is under Ca^2+^ regulation, remains unknown. In axons of living neurons, we found KCl-induced depolarisation generated Ca²⁺ "hot spots" with persistently elevated Ca²⁺, leading to focal aggregation of ANXA7 in them (Fig. 4A-J). Unlike ANXA11, which tethers RNPs to lysosomes in a Ca^2+^-dependent manner ^11^, ANXA7 molecules respond to Ca^2+^ surges by rapidly forming droplets that are recruited to the ruptured plasma membrane (PM). This recruitment facilitates the ESCRT III-dependent repair process in cancer cells ^45^. Our observation of Ca^2+^-induced ANXA7 focal aggregation aligns with these findings, showing focal aggregation near the PM in axons. However, further research is needed to explore the specific functions of these Ca^2+^-induced ANXA7 aggregates in axons.

We identified that Ca^2+^ elevation enhanced the LLPS of ANXA7, leading to the formation of ANXA7 droplets (Fig. 3D-E), which engaged the TIA1 into themselves (Fig. 3G-I). Significantly, employing a confocal microscopy-based *in vitro* approach, we found that Ca^2+^ causes the detachment of small TIA1/ANXA7 droplets from the DIC1B coated on beads, and forming large aggregate-like sediments at the bottom of the dish (Fig. 3J-O). Consistently, by experiments in neurons, axon trafficking is found to be dramatically inhibited, leading to axonal aggregation of both TIA1 and ANXA7 (Fig. 4K-P). However, since the precise domain involved in the interaction between ANXA7 and DIC1B is not yet identified, the exact mechanism behind their detachment remains unclear. Moreover, knockdown of endogenous ANXA7 leads to aggregation of TIA1 in axons (Fig. 5A-C; Fig. 5K-L), which in turn causes the axonopathy in CST of mice spinal cord and neurodegeneration (Fig. 6D-H). This finding is consistent with the established role of TIA1 granules in promoting degeneration in Tau P301S mice ^27,28^ and their regeneration-suppressing effects in both worm ^24^ and rodent neurons ^34^, supporting the notion that TIA1 aggregation is pathogenic for axon health. Therefore, we detected not only a novel direct axonal trafficking mechanism of RNP, but also a Ca^2+^ overload-triggered pathological mechanism underlying axonopathy.

### Up-regulation of ANXA7 represses TIA1 aggregates in axons

In living axons, we found that increasing the level of ANXA7 enhances the dynamics of axonal TIA1 granules. Conversely, knocking down ANXA7 results in more immobile and condensed TIA1 droplets (Fig. 5D-J), underscoring the crucial role of ANXA7 in controlling the LLPS dynamics of TIA1 granules in axons. Consistently, knockdown endogenous ANXA7 leads to pathological TIA1 aggregates, which, in turn, cause axonopathy and neurodegeneration both *in vitro* (Fig. 5K-P) and *in vivo* (Fig. 6D-N). Protein aggregation within axons is regarded as a pathogenic reason for neurodegenerative diseases, including ALS, FTD, and WDM ^21,69,70^. Specifically, mutations in *TIA1* have been linked to WDM ^26^, FTD, ALS ^25,71-73^, and multisystem proteinopathy (MSP) ^74^. Noticebly, all of these pathological *TIA1* mutations are located in its PrLD, which facilitates the formation of toxic TIA1 aggregates ^25,74,75^. Additionally, LLPS of wild-type TIA1 has been found to promote the phase separation and toxic oligomerisation of tau, exacerbating tauopathies ^28^. Reducing TIA1 levels can inhibit the accumulation of tau oligomers and improve neuronal survival in tauopathy mouse models ^27,28^, further underscoring the pathogenic role of TIA1 aggregates in these neurodegenerative diseases.

Therefore, our finding that the overexpression of ANXA7 alleviates the formation of TIA1 aggregates is highly promising. This suggests that boosting ANXA7 level could represent a potentially effective therapeutic strategy for treating TIA1 aggregation-related neurodegenerative diseases.

Our research has identified that most TIA1 granules move in a retrograde direction toward the cell body. However, the precise composition, function, and fate of these retrograde TIA1-containing RNPs remain largely unexplored. Previous research has established that TIA1 binds to the 3’UTR of mRNA and represses its translation in several types of cells ^29-32^. In the nervous system, TIA1 has been found to suppress the expression of mRNAs related to neurodevelopment ^33^. Following axon injury, TIA1 suppresses the translation of regeneration-associated mRNAs, such as *Neuritin1 (Nrn1)* and *Importin β1 (Impβ1)* ^34^. Yet, the exact mRNA cargoes of TIA1 in axons remain unknown. Investigating whether a distinct pool of mRNAs and RBPs specifically recruited to the retrograde flux of TIA1 granules could provide valuable insights into their function and ultimate destination. Overall, exploring the retrograde transport of RNPs could yield significant insights into the biology and pathology of long-extending neurons.

## Methods

### Antibodies and DNA constructs

Primary antibodies: anti-TIA1 (#sc-166247, Santa Cruz), anti-TIA1 (#12133-2-AP, Proteintech), anti-SQSTM1/p62 (#A11247, ABclonal), anti-G3BP1 (#sc-365338, Santa Cruz), anti-Rab5 (#3547, Cell Signaling Technology), anti-LC3B (#83506, Cell Signaling Technology), anti-LAMP1 (#ab13523, Abcam), anti-DYNC1I1 (#13808-1-AP, Proteintech), anti-ANXA7 (#10154-2-AP, Proteintech), anti-GAPDH (#10494-1-AP, Proteintech), anti-HA (#3724, Cell Signaling Technology), anti-Myc (#60003-2-Ig, Proteintech), anti-Myc (#16286-1-AP, Proteintech), anti-Flag (#20543-1-AP, Proteintech), anti-TDP-43 (#89789, Cell Signaling Technology), anti-Iba1 (#019-19741, FUJIFILM).

Secondary antibodies: anti-mouse IgG for IP (HRP) (#ab131368; Abcam), HRP-labeled goat anti-mouse or rabbit IgG (H+L) (#A0216, #A0208; Beyotime), Veriblot for IP detection (#ab131366; Abcam). All fluorescent-dye conjugated secondary antibodies were from Invitrogen.

ANXA7 was cloned from rat brain cDNA library using primer pair (forward, 5’-CGCCTCGAGCTTAAGTATGTCATACCCAGGCTACCC-3’; reverse, 5’-CACTATAGAATAGGGCCCTTCACTGGCCAACGATGGC-3’). All primers were synthesised from Tsingke (China). The shRNAs targeting rats or mice were designed using the ThermoFisher BLOCK-iT RNAi Designer or Sigma-Aldrich website, respectively, and synthesised by GENEWIZ (China). siRNAs were synthesised by Genomeditech (China). All sequences are listed in the Supplementary Table 4.

The DNA encoding TIA1 was provided by Prof. Yichang Jia (Tsinghua University). The DNA encoding DIC1B-mRFP was provided by Prof. K. Kevin Pfister (University of Virginia). Lifeact-GFP was provided by Prof. Roland Wedlich Soldner (MPI Biochemistry). The DNA encoding Cry2 was provided by Prof. Hanhui Ma (ShanghaiTech University). GCaMP6f was purchased from Addgene (#40755). pAAV-hSyn-GFP, pHelper, and pPHP.S were gifts from Prof. Zhenge Luo (ShanghaiTech University).

### Primary neuronal culture and transfection

Hippocampal or cortical tissues were derived from embryonic day 18 (E18) Sprague– Dawley rat brains, following relevant guidelines and regulations as approved by the Animal Ethics Committees of ShanghaiTech University (approval number: 20230217002). Then neurons were dissociated, suspended in plating medium (DMEM with 10% FBS, 10% F-12 and 1% Penicillin-Streptomycin), and seeded on Poly-L-Lysine coated glass coverslip, 29 mm glass bottom dish (#D29-20-1.5-N, Cellvis) at 3.2 × 10^4^ cells/cm^2^, or into polydimethylsiloxane (PDMS) microfluidic device at 2 × 10^5^ cells per reservoir, as previously described ^36,76,77^. Plating medium was half changed to maintain medium (Neurobasal Mediumwith 2% B27 and 1% L-GlutaMax) on DIV1, andon DIV3-4, 10 µM 5-fluoro-2’-deoxyuridine (FDU) was added to suppress non-neuronal cell growth. For hippocampal neurons, DIV6 neurons were transfected with 1-2 µg indicated plasmids using Lipofectamine 2000. Cortical neurons were electroporated using Nucleofector 2b (Lonza) before seeding and harvested on DIV8-11 for western blot analysis.

### Analysis of granule trafficking in live axons

To fluorescently label RNPs in axons, DIV6 rat hippocampal neurons were cultured in 29 mm glass-bottom dishes and co-transfected with 0.2 nmol of CY5-UTP (#B8333, APExBIO) for total RNA labelling and 1 µg of GFP-TIA1. Live-imagingwas then conducted on DIV8 by replacing the medium with imaging buffer (15 mM HEPES, 145 mM NaCl, 5.6 mM KCl, 2.2 mM CaCl₂, 0.5 mM MgCl₂, 5.6 mM D-glucose, 0.5 mM ascorbic acid, 0.1% BSA, pH 7.4). Time-lapse confocal images were acquired using the Nikon TI2-E inverted microscope equipped with a Yokogawa spinning confocal disc head and a 60 × 1.4 NA oil objective, with 1-5 seconds interval. Acquired time stacks were analysed in ImageJ (v2.3.0/1.53f, NIH). Kymographs of RNP movement were generated using the Multi-Kymograph plugin. Directions were assigned based on relative location to the soma.

For TIA1 granule axon trafficking analysis, rat hippocampal neurons at DIV6 were cultured in microfluidic device or glass-bottom dish and transfected with TIA1-GFP/mCherry or other indicated plasmids. DIV8 cells were imaged under the same conditions with 5 to to 22-second interval and analysed using ImageJ, as previously described ^77^. Briefly, axonal TIA1 granules were traced using the Trackmate plugin (v7.11) with an estimated object diameter of 0.8 µm. Granules used for statistical analysis were filtered based on track duration (over 2 frames) and track speed (0-2 µm/s). The mean speeds of the tracks were exported, and directions were assigned as described above.

For co-trafficking analysis of TIA1 granules, DIV6 rat hippocampal neurons cultured in microfluidic devices were either co-transfected with GFP-TIA1 and DIC1B-mRFP or co-labelled on DIV8-9 using a pulse-chase labelling assay as previously described ^77^. Briefly, the axon terminal chamber was incubated with an imaging buffer containing either 100 nM BoNT/A-Hc-Atto647N, 50 ng/ml Alexa Fluor 647-conjugated recombinant CTB (#C34778, Invitrogen), or 50 nM Lysotracker Red (#40739ES50, Yeasen) for 10 minutes at 37°C. Axons within the channels and terminal chambers were live-imaged under the same conditions, with 4-22 seconds interval. Total number of TIA1 granules and their co-trafficking proportions with dynein or membranous markers were manually counted in kymographs in ImageJ.

### 1,6-Hex-induced granule diffusion analysis

To assess the response of TIA1 granules to 1,6-Hex, DIV6 rat hippocampal neurons were transfected with TIA1-mCherry. Live imaging was performed using a confocal microscope under the same conditions above on DIV12. Time-lapse images were captured before and after 1,6-Hex (#240117, Sigma-Aldrich) treatment with 11-16 seconds interval and analysed using ImageJ. Kymographs of axonal TIA1 granules were generated and the TIA1 heterogeneity index was calculated as detailed in sFig. 1L.

### OptoDroplet assay

Hippocampal neurons were cultured in glass-bottom dishes and transfected with Opto-Control, Opto-TIA1, and indicated plasmids on DIV6. Live-cell imaging was performed on DIV9 using previous confocal microscope with a 40× 1.3 NA oil objective. Neurons were exposed to combined laser excitation at 561 nm for mCherry imaging and 488 nm for blue light activation of Cry2. Time-lapse images were continuously acquired over 20 minute span with 5-12 seconds interval, and approximately 120 seconds were sufficient for Opto-TIA1 granule formation. The acquired time-stacks were analysed in ImageJ. Heterogeneity indexes were calculated as described in sFig. 1L. The first frame was designated as the "Before" state, while the first frame after 120 seconds light activation as the "After" state. Axonal granule number was manually assessed and normalised to axon length. Following 120 seconds of blue light activation, time-lapse images were analysed using the TrackMate plugin to trace and analyse the movement of axonal Opto-TIA1 granules, following the protocol described above.

### Analysis of high K⁺-induced intracellular Ca^2+^ responses in live neurons

On DIV12-13, rat hippocampal neurons expressing the Ca²⁺ sensor GCaMP6f were subjected to high K⁺ stimulation following previously established protocols ^38^. Briefly, the culture medium was replaced with a warm high K⁺ buffer (same as the imaging buffer except that it contained 95 mM NaCl and 56 mM KCl), whereas control neurons were treated with an imaging buffer. Imaging was performed as above using a confocal microscope with a 60× 1.4 NA oil objective, and time-lapse images were captured continuously before and immediately after the medium was replaced, with 12-22 seconds intervals. The resulting time stacks were analysed using ImageJ, adhering to previously outlined analysis steps.

### Lattice SIM and analysis

To examine the co-localisation of endogenous TIA1 with ANXA7 or DIC1B in axons, rat hippocampal neurons transfected with indicated plasmids on DIV6 were fixed and stained on DIV12. Imaging was conducted using Lattice SIM on a ZEISS Elyra 7 microscope with a 63× 1.4 NA oil objective, utilising a grid size of 27.5 μm with 12 rotations. Raw SIM images were processed with Fourier transformation in Zen software (version 16.0.13.306, ZEN 3.0 SR black edition; Zeiss), followed by the application of a sharpness filter and fast fit advanced filter. The processed 3D-SIM images were analysed in ImageJ. Colocalisation rates between the two channels were calculated as Pearson’s correlation coefficients using the JACoP plugin.

### Protein expression and purification

His-TEV-Flag-DIC1B and His-TEV-Myc-ANXA7 proteins were expressed in *E. coli*. BL21(DE3) cells. Cultures were induced at an OD₆₀₀ of 0.6-0.7 with IPTG (1 mM and 0.5 mM, respectively) at 16°C for 16 hours. The cells were resuspended in lysis buffer (50 mM phosphate buffer, 300 mM NaCl, 50 mM L-arginine, 2 mM MgCl₂, 2 mM imidazole, pH 7.0) with 0.2 mM PMSF, 2 mM DTT, 1 mM protease inhibitor, and 20 U/mL DNase I. After sonication, soluble proteins were separated by centrifugation at 18,000g for 30 minutes. The soluble fraction was incubated with Ni Sepharose 6 Fast Flow resin (#17531802, Cytiva), washed with wash buffer (50 mM phosphate buffer, 300 mM NaCl, 10% glycerin, 2 mM DTT, 50 mM imidazole, pH 7.0), and eluted with elution buffer (50 mM phosphate buffer, 300 mM NaCl, 0.5 M L-arginine, 200 mM imidazole, pH 7.0, 2 mM DTT). The eluted fractions were dialysed against TEV cleavage buffer (50 mM phosphate buffer, 300 mM NaCl, 0.5 M L-arginine) to reduce imidazole concentration to below 0.2 mM. Protein concentration was estimated by SDS-PAGE using BSA standards. TEV protease was added to the protein solution at a 1:30 enzyme-to-protein mass ratio and incubated at 4°C for 24 hours. Following cleavage, the mixture was treated with Ni Sepharose 6 Fast Flow resin to remove the tag, and the target protein was collected from the flow-through. The purified protein was concentrated and stored in aliquots at -80°C.

The expression and purification of GST-TEV-TIA1 followed a similar protocol, substituting Glutathione Sepharose 4 Fast Flow resin (#17513202, Cytiva) for affinity purification. Imidazole was excluded from all buffers, and the elution buffer contained 10 mM reduced glutathione. TEV protease was used to cleave the GST tag from TIA1, and the resulting proteins were purified using size-exclusion chromatography on a HiLoad 16/600 Superdex 75 pg column (#28989333, Cytiva) with an ÄKTA Pure system (Cytiva).

### Co-IP, in vitro pull-down and western blot

For the Co-IP assay, HEK293T cells were harvested 48 hours post-transfection, while primary cortical neurons were collected 11 days post-electroporation. Cells were washed with cold PBS and lysed in NP40 lysis buffer (50 mM Tris-HCl, pH 8.0, 150 mM NaCl, 5 mM MgCl₂, 0.5% NP40) containing protease inhibitors. Lysates were then centrifuged at 21,400g for 10 minutes at 4°C, and the supernatants were collected for IP. IP was performed using Anti-Flag M2 Affinity beads (#A2220, Sigma-Aldrich) or Anti-HA magnetic beads (#B26202, Bimake) with a 3-hour incubation at 4°C. Beads were subsequently washed with NP40 wash buffer (50 mM Tris-HCl, pH 8.0, 300 mM NaCl, 5 mM MgCl₂, 0.1% NP40), and the bound proteins were eluted for western blot detection.

For the *in vitro* pull-down assay, 2 μg Flag-DIC1B was incubated with Anti-Flag magnetic beads (#B26102, Selleck) in NP40 lysis buffer with 0.5 M L-arginine at 4°C for 1 hour. Following this, 2 μg each of Myc-ANXA7 and TIA1 were added and the incubation was continued at 4°C for an additional hour. After incubation, the supernatants were removed, and the beads were washed five times with NP40 wash buffer with 0.5 M L-arginine. The beads were then mixed with 1× SDS loading buffer. For experiments involving Ca²⁺, 1 mM CaCl₂ was added to both the NP40 lysis and wash buffers. Interactions were quantitatively analysed by western blot.

Western blot samples were separated on 8% or 10% Tris-glycine polyacrylamide SDS-PAGE and transferred to PVDF membranes. Membranes were then blocked with 5% non-fat milk in TBST (0.05% Tween) for 1 hour at room temperature, followed by overnight incubation with primary antibodies at 4°C. The membranes were washed and then incubated with secondary antibodies for 1 hour at room temperature. Primary antibodies were diluted at 1:2000, while secondary antibodies were diluted at 1:5000, except for Veriblot which was diluted at 1:1000. Blots were detected immediately using the Amersham Imager 680 (Cytiva) or the Touch Imager (e-BLOT Life Science).

### Proteomic analysis of brain interactomes

Cortical tissues from P14 rats were homogenised on ice in homogenisation buffer (0.32 M sucrose, 10 mM HEPES, pH 7.4) and lysed in four volumes of RIPA buffer (50 mM Tris-HCl, 150 mM NaCl, 1% NP40, 0.25% sodium deoxycholate, pH 7.4) with protease inhibitors. The lysates were centrifuged at 15,000g for 40 minutes at 4°C to remove debris. Supernatants were quantified using the Bradford assay and subsequently pre-cleared with glutathione resin. The purified proteins were blocked with 1% BSA for 1 hour at 4°C, then added to the brain lysates and incubated at 4°C for an additional 2 hours. After this incubation, glutathione resin beads were introduced and incubated under the same conditions for another 2 hours. The beads were then pelleted by centrifugation at 1000g for 5 minutes and washed four times with NP40 wash buffer. Western blot analysis was subsequently performed as previously described.

For proteomic analysis following the GST pull-down, the prepared samples were processed for mass spectrometry. Gel strips were cut into 1.5 mm pieces and washed, then decolorised using a 25 mM NH₄HCO₃/acetonitrile (1:1) solution, dehydrated with acetonitrile, and vacuum-dried. The proteins were reduced with 10 mM DTT at 56°C for 1 hour and then alkylated with 25 mM Iodoacetamide (IAM) for 45 minutes in the dark. Following sequential washes with 25 mM NH₄HCO₃, a 25 mM NH₄HCO₃/acetonitrile (1:1) solution, and acetonitrile, the samples were vacuum-dried again. Proteins were digested by adding an enzyme at a 50:1 protein-to-enzyme ratio. The samples were incubated at 4°C for 20 minutes and then at 37°C overnight. The resulting peptides were extracted with 50% acetonitrile/0.5% formic acid, combined, and vacuum-dried. The peptides were redissolved in 0.1% formic acid and desalted using Stage-Tips.

Proteomic analysis was conducted using a Q Exactive HF-X mass spectrometer (Thermo Fisher Scientific). The mass spectrometry (MS) data were analysed using Proteome Discoverer (version PD2.2) and normalised by total protein intensity. Seq-k-nearest neighbor (Seq-Knn) imputation was applied for missing values using the ‘Wu Kong’ platform (https://www.omicsolution.com/wkomics/wkold/). Differential analysis between GST control and GST-TIA1/GST-ANXA7 was based on a fold-change (FC) ≥ 2.3 (log2FC ≥ 1.2) and a P-value ≤ 0.05 (-log10P ≥ 1.3). GO and KEGG pathway analyses were performed using Metascape ^78^, and enrichment dot bubble plots were generated on https://www.bioinformatics.com.cn.

### FRAP assay

48 hours after being transfected with GFP-TIA1 plasmids on DIV6, rat hippocampal neurons in glass-bottom dishes were placed on the previously described confocal microscope with a 60× 1.4 NA oil objective for live-imaging, with 1 second interval. GFP signals were bleached using a 488-nm laser set at 90% intensity for 100 ms, following the acquisition of six pre-bleach images. Then, the neurons were allowed to recover and recorded for 5 minutes after photobleaching.

Time-lapse images were processed and analysed using Nikon Elements AR, and time measurement results were exported. FRAP efficiency (E) was calculated using the equation (1):

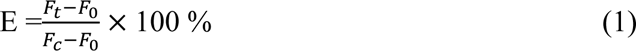

where F_t_ is the intensity at time t, F_0_ is the intensity immediately after photobleaching, and F_c_ is the intensity before photobleaching (corrected by Exponential One phase decay in GraphPad).

### FLIM-FRET assay

Primary hippocampal neurons were transfected with the indicated plasmids (Donor: GFP-TIA1, Acceptor: DIC1B-mRFP) on DIV6. On DIV9, neurons were imaged in prewarmed imaging buffer using a Leica STELLARIS 8 FALCON microscope with a 63× 1.4 NA oil objective. The tunable white light laser was set to 489 nm excitation at an 80 MHz frequency. Emission from 494-540 nm was collected using a HyD X1 detector, and laser power was adjusted to achieve approximately 1 photon per laser pulse, following the published method ^79^. Confocal settings included a 512×512 pixel resolution with a 4.0 optical zoom, resulting in a 0.09 μm pixel size. FLIM images were processed using LAS X 4.4.0.24861 software (Leica Microsystems). Lifetime decay curves were fitted with an n-Exponential Reconvolution model, selecting the number of components based on χ² values closest to 1. FLIM images were analysed using ImageJ.

FRET efficiency (*E*_FRET_) was calculated in equation (2):

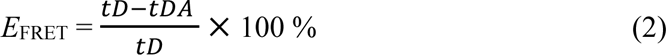

where t_DA_ is the donor lifetime (GFP-TIA1) in the presence of the acceptor (DIC1B-mRFP), and t_D_ is the donor lifetime without the acceptor.

### In vitro phase separation assays

For the Ca²-induced sedimentation assay, purified Myc-ANXA7 and TIA1 proteins were desalted using SpinDesalt columns (#SEC02301, Smart-Lifesciences), and their concentrations were determined. A 100 μL reaction mixture containing 5 μM protein in LLPS buffer (50 mM HEPES, 50 mM NaCl, 2 mM DTT) was prepared, followed by the addition of CaCl₂ (0-2 mM). The mixture was incubated at 4°C for 30 minutes and then centrifuged at 14,000g for 20 minutes at 25°C. A 10 μL aliquot from the supernatant (S) and the condensed phase (P) was subjected to SDS-PAGE and stained with Coomassie Blue, and phase separation was quantified by measuring the band intensities.

For fluorescently labeled phase separation assays, fluorescent dyes (Atto 647N NHS ester (#05316-1MG-F, Sigma-Aldrich), Vari Fluor 568 SE (#HY-D1799, MedChemExpress), or iFluor 488 succinimidyl ester (#1023, AAT Bioquest)) were conjugated to purified proteins (2 mg/ml) in bicarbonate buffer (0.1 M, pH 8.3) at a dye-to-protein ratio of 10:1. After a 1-hour incubation at room temperature, labeled proteins were desalted using SpinDesalt columns, and their concentrations were measured. Proteins were then mixed in phase separation buffer (10 mM HEPES, 150 mM NaCl, 0.1 mM EDTA, 2 mM DTT, pH 7.4), with PEG8000 (#P8260, Solarbio) or Ca^2+^ in some cases. Samples were injected into homemade imaging chambers, consisting of a coverslip and glass slide held together by double-sided tape. After a 15-minute incubation at room temperature, the chamber was imaged using the Leica Thunder Imager (DMi8) with HC PL APO 63 × 1.40 NA oil objective (Figure 4F), Nikon TI2-E inverted microscope equipped with a Yokogawa spinning confocal disc head (CSU-W1) and a 60 × 1.4 NA oil objective (Figure S4A and S4E), or Olympus SpinSR with a UPLXAPO 20 × objective (Figure 4D, 4G and Figure S4F).

For microscopy-based phase separation assay, to generate the DIC-647 coated beads, 20 µl of Anti-Flag M2 Affinity beads were incubated with Atto-647-labbled Flag-DIC1B (DIC-647, 5 µM) at 4°C for 1h and washed by 200 µl imaging pull-down buffer (25 mM Tris-HCl pH 7.5, 150 mM NaCl, 1 mM DTT). The DIC-647-coated beads were centrifuged at 1500g for 2 minutes at 4°C and resuspended in 10 µL of imaging buffer. Then, 2 µL of resuspended beads were added to 20 µL of imaging mixture containing 5 µM ANXA7-568 and 5 µM TIA1-488. After a 30-minute dark incubation, the 20 µL imaging mixture was placed in the center of a slide with double-sided tape, and a coverslip was placed on top. The slide was inverted and immediately imaged using an Olympus SpinSR with a UPLXAPO 20× objective to obtain 3D stacks of the coated beads. All experiments were conducted in darkness to minimise bleaching.

### IF staining and 3D rendering analysis

Primary rat hippocampal neurons at DIV9 or DIV12 were fixed in 4% PFA and 4% sucrose in PBS for 30 minutes at room temperature. After blocking in antibody diluting buffer (0.1% saponin, 1% BSA, 0.2% gelatin in PBS) for 1 hour at room temperature, cells were incubated with primary antibodies (1:500) overnight at 4℃, followed by secondary antibodies (1:5000) for 1 hour in the dark at room temperature. DAPI in PBS was added for 10 minutes at room temperature, and cells were mounted with Fluoroshield mounting medium. For PI staining, PI (#P1304MP, Invitrogen) was added after DAPI incubation according to the product manual.

Z-stack images were captured using a Nikon TI2-E inverted microscope equipped with a Yokogawa spinning confocal disk head (CSU-W1 2 camera) with a 60 × 1.4 NA oil objective. Images were analysed in ImageJ. Co-localisation rates between two channels were measured using Pearson’s coefficient with the JACoP plugin. For granule detection, axon shafts with lengths ≥ 50 µm were selected and straightened. Granules were identified using the "Analyse Particles" function with a circularity > 0.6 and area between 0.05-20 μm². Detected granule sizes were exported, and granule numbers were normalised to axon length.

To analyse the axonal co-localisation of TIA1 with G3BP1 or p62, the 3D stacks of confocal images were deconvoluted using 40 cycles of Huygens Professional software (v18.10, Scientific Volume Imaging) and imported into Imaris software (v9.7.2, Bitplane) for morphology fitting using the “surface” function. For precise 3D renderings, "Surface Grain Size" was set at 0.100 µm and "Diameter of Largest Sphere" at 1.00 µm for neurons co-labeled with TIA1 and G3BP1, while for neurons co-labeled with TIA1 and p62, parameters were 0.00100 µm and 1.00 µm respectively.

### AAV packaging and ICV injection

The pAAV-U6-hSyn-GFP plasmid was constructed by cloning the U6 promoter into the pAAV-hSyn-GFP vector using specific primers (forward: 5’-CGGCCGCACGCGTGTGTGAGGGCCTATTTCCCATGAT-3’; reverse: 5’-CAGGGCCCTCTGCAGTCTAGAGGTGTTTCGTCCTTTCCAC-3’). shRNA targeting mouse ANXA7 (shANXA7-3# and shANXA7-4#; sTab. 4) was synthesized and inserted into the pAAV-U6-hSyn-GFP plasmid. AAV particles were packaged into serotype 9 capsids and purified as previously described ^77^. Briefly, target plasmid and AAV helper plasmid (pPHP.s) were co-transfected into HEK293T cells using PEI. After 72 hours, cells were harvested, lysed with buffer (150 mM NaCl, 20 mM Tris-HCl, pH 8.0), treated with Benzonase for 45 minutes, and centrifuged to collect supernatant. AAV particles were purified by OptiPrep density gradient ultracentrifugation (40% fraction) and titrated by qPCR targeting the GFP region.

All animal procedures were conducted under the ethical guidelines of the Institutional Animal Care and Use Committee of ShanghaiTech University (approval number: 20230217002). C57BL/6J mice were housed on a 12-h light-dark cycle. Both male and female pups at postnatal day 1 were cryoanesthetized and AAV were bilaterally injected into the cerebral ventricles using Drummond™ PCR Micropipets, pulled with a P-97 Flaming/Brown micropipette puller. Each ventricle received 1 μl of virus mixed with Fast Green for visualisation.

### Rotarod test

Motor coordination of 8-week-old mice was assessed using a Rotarod machine (Ugo Basile, Model 47650). Mice were trained over three days (P56-P58) and tested on the fourth day (P59). Training involved constant speed at 4 rpm on P56, 4-20 rpm acceleration over 5 minutes on P57, and 4-40 rpm on P58. On P59, mice performed three trials at 4-40 rpm acceleration over 5 minutes, with a 20 minutes rest between trials. The average time to fall off from three trials was recorded for each mouse.

### Tissue sectioning, staining, imaging, and analysis

P60 mice were deeply anesthetised with isoflurane and perfused with 0.9% saline followed by 4% PFA in PBS. The brain and spinal cord were dissected, post-fixed in 4% PFA at 4°C overnight, and then dehydrated in 30% sucrose in PBS at 4°C until sinking. Tissues were embedded in OCT (#4583, Sakura) and sectioned using a cryostat microtome, with brains coronally at 40 μm and spinal cords longitudinally at 25 μm. For IF staining, sections were permeabilised with 1% SDS in PBS for 4 minutes, blocked with 1% BSA in PBS for 1 h at room temperature, and incubated with primary antibodies (1:500) overnight at 4°C. Then, they were incubated with secondary antibodies (1:5000) for 3 h at room temperature, and stained with DAPI for 10 minutes. Sections were mounted with Fluoroshield mounting medium. All steps were conducted in the dark. Images were captured using an Olympus VS120-S6-W slide scanner with a 20× 0.5 NA objective. Focused imaging was performed using a Nikon TI2-E inverted microscope equipped with a Yokogawa spinning confocal disk head (CSU-W1 2 camera) with a 60× 1.4 NA oil objective.

For axon morphological analysis, GFP-labeled axons were quantified by measuring the width-to-height ratio of bounding rectangles for each signal using ImageJ. Axons with width-to-height ratios 0.5-2 were classified as degenerating fragments. The total areas of infected and degenerating axons were used to calculate the percentage of degeneration. For the percentage of active microglia in the motor cortex, Iba1^+^ microglia and DAPI^+^ cells were quantified using the "Analyse Particles" function, with an area threshold >5.27 μm² and normalised to the number of DAPI^+^ cells.

### Statistical information

All data were illustrated and analysed using GraphPad Prism (v8.3.0). For live-cell experiments, results were obtained from more than two independent neuron preparations, whereas fixed samples were analysed from at least three independent preparations. Results are presented as mean ± SEM. Data sets that followed a normal distribution between two groups were analysed using a two-tailed unpaired *t*-test to determine statistical significance. For paired data comparisons, a two-tailed paired *t*-test was used. One-way ANOVA was applied for comparing two or more groups against a control. Statistical significance was defined as a *p*-value of less than 0.05. For sFig. 1B; sFig. 1N; Fig. 5H and Fig. 5N, outliers were identified and excluded using the ROUT method with Q = 1%. Sample size adequacy was determined based on preliminary data or through discussion. ROI selection was randomised to avoid bias. Data collection and analysis were performed by independent operators who were blinded to the experimental conditions.

### Data availability statement

All data reported in this paper will be shared by the corresponding author upon request. This paper does not report the original code. Any additional information required to reanalyse the data reported in this paper is available from the corresponding author upon request.

## Acknowledgements

We thank Professor F.A. Meunier, Professor Lei Li, on their constructive comments, and thank Dr Xiaoming Li, Dr. Zhaomei Shi and Dr Ziwei Yang for her/his expert technical assistance. This work was supported by the National Natural Science Foundation of China (32271001 and 31871036 to T. Wang). Y. Chu acknowledges the support from the National Natural Science Foundation of China (32100777). Y. Liu would like to thank the Double First-Class Initiative Fund of ShanghaiTech University (SYLPOC0022022, SYLDX0302022).

## Author contributions

Conceptualisation: T. Wang, Y. Feng; Methodology: Y. Feng, T. Luan, Z. Zhang, Y. Chu, X. Pan, J. Li; Investigation: Y. Feng, T. Luan, Y. Chu, W. Wang, S. Wan; Visualization: T. Wang, Y. Feng, T. Luan, W. Wang, S. Wan; Tools: J. Li, Y. Liu; Supervision: T. Wang, Y. Liu; Writing—original draft: T. Wang, Y. Feng, T. Luan; Writing—review & editing: T. Wang, Y. Feng, T. Luan;

## Materials & Correspondence

Correspondence and material requests could be addressed to the corresponding author Tong Wang.

---

**Supplementary Figure 1.**
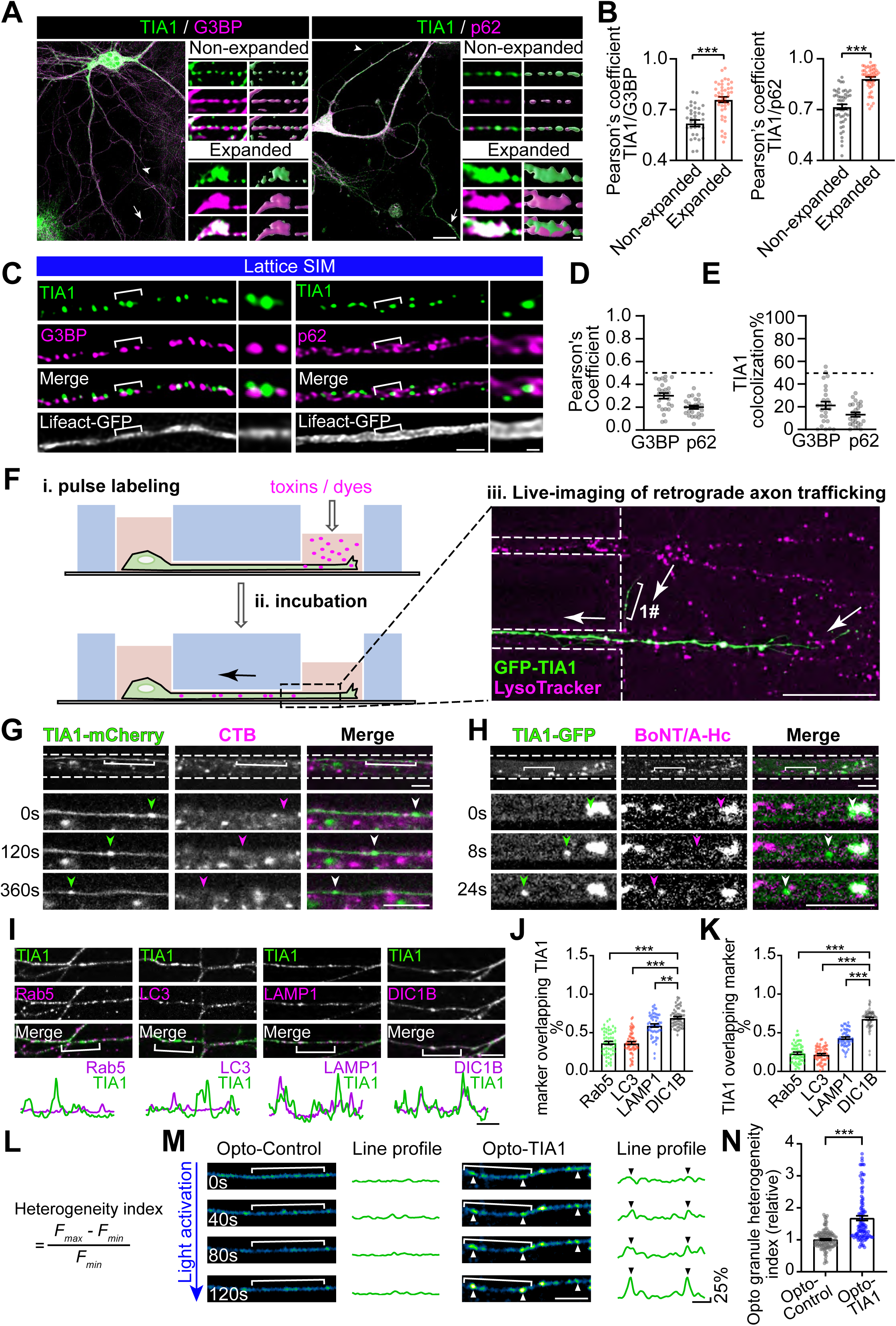
TIA1 granules undergo retrograde trafficking in axons. **(A)** Representative images showing the distribution of endogenous TIA1 with G3BP or SQSTM1/p62 in DIV9 cultured rat hippocampal neurons, with Imaris rendering in the right panels. Arrowheads indicate unexpanded axon segments; arrows indicate expanded segments. Scale bar = 20 µm (left), 1 µm (right). **(B)** Pearson’s coefficient of TIA1 with G3BP or SQSTM1/p62 in unexpanded and expanded axon segments (For G3BP: n = 36, 43; for p62: n = 48, 47). **(C)** Lattice SIM images showing the distribution of endogenous TIA1 with G3BP or SQSTM1/p62 in axons. Scale bar = 1 µm (left), 0.2 µm (right). **(D)** Quantification of Pearson’s coefficient for TIA1 co-localization with G3BP or p62 in axons. **(E)** Ratio of TIA1 co-localized with G3BP or p62 in axons (n = 25). **(F)** Schematic diagram of the pulse-chase labelling assay to specifically label retrograde membranous axonal organelles in neurons cultured in a microfluidic device. See also the Methods for details. Scale bar = 50 µm. **(G-H)** Time-lapse images showing TIA1 granules trafficking with CTB **(G)** or BoNT/A-Hc **(H)**. Arrowheads indicate moving TIA1 granules. Scale bar = 10 µm. **(I)** Representative confocal images of endogenous TIA1 with organelle markers (Rab5 for endosomes, LC3 for autophagosomes, LAMP1 for lysosomes, DIC1B for dynein) in axons, with intensity profiles shown below. Scale bar = 10 µm (top), 5 µm (bottom). **(J-K)** Quantification of **(I)**, with **(J)** showing the ratio of TIA1 co-localized with the indicated markers, and **(K)** showing the ratio of the markers co-localized with TIA1 (n = 56, 56, 53, 56). **(L)** Equation of the heterogeneity index. **(M)** Time-lapse images showing intensity changes induced by blue light in axons expressing Opto-Control or Opto-TIA1. Intensity profiles in the bracketed regions are shown on the right. Arrowheads indicate light-induced Opto-TIA1 granules. Scale bar = 10 µm (left), 5 µm (right); y-axis = 25%. **(N)** Quantification of **(M)** showing heterogeneity changes induced by blue light exposure for Opto-Control and Opto-TIA1 (n = 121, 114). Data represent mean ±SEM; in **(B, N)** two-tailed unpaired *t*-test; in **(J, K)** one-way ANOVA. \*\**p*<0.01, \*\*\**p*<0.001.

**Supplementary Figure 2.**
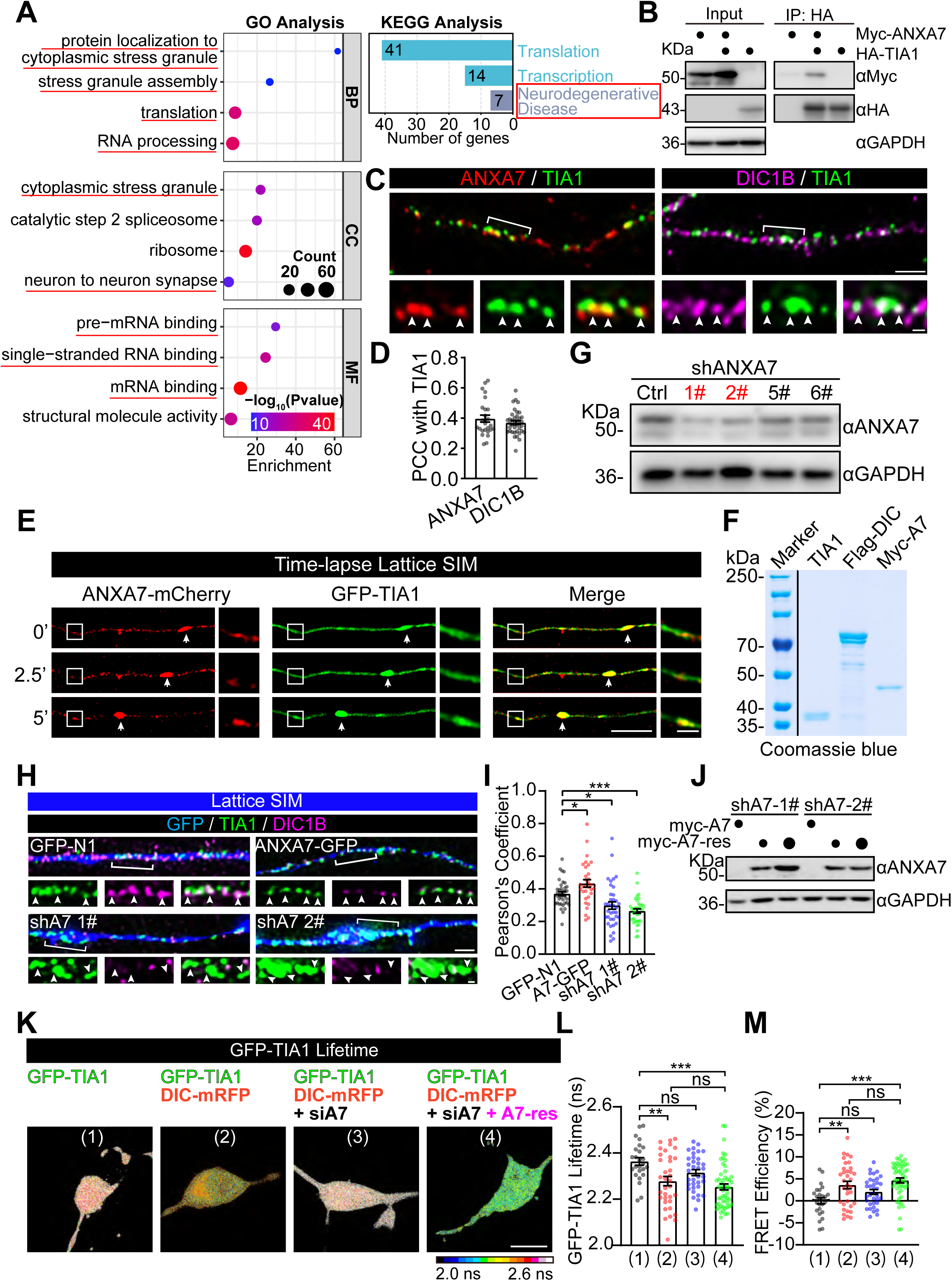
ANXA7 promotes the recruitment of TIA1 granules to dynein. **(A)** GO and KEGG enrichment pathway analysis for proteins pulled down by GST-TIA1 from P14 rat brain lysates. **(B)** Co-IP of Myc-ANXA7 and HA-TIA1 co-expressed in HEK293T cells, showing interaction. **(C)** In DIV12 rat hippocampal neurons, 3D-Lattice SIM images of TIA1 (green) and ANXA7 (red) or DIC1B (magenta) along axons. Arrowheads indicate co-localization. Scale bars = 1 μm and 200 nm. **(D)** Pearson’s coefficient for co-localization between TIA1 and ANXA7 or DIC1B from **(C),** (n = 26, 38). **(E)** The arrows in time-lapse lattice SIM images showing retrograde co-trafficking of granules in DIV13 hippocampal neurons. One newly formed TIA1/ANXA7 granule (boxed) are amplified in the right panels. Scale bars = 5 μm and 1 μm. **(F)** Coomassie blue staining of an SDS-PAGE gel showing the purity of TIA1, Flag-DIC1B (Flag-DIC) and Myc-ANXA7 proteins expressed in *E. Coli*. **(G)** Western blot analysis validating the efficiency of shRNA-mediated knockdown targeting the rats endogenous ANXA7 (shANXA7) in cultured rat cortical neurons. shRNA sequences of 1# and 2# are available in Methods. **(H)** Representative 3D lattice-SIM images showing the distribution of endogenous TIA1 (green) and DIC1B (magenta) in axons of DIV12 cultured rat hippocampal neurons under conditions of endogenous ANXA7 knockdown (shA7 1# and 2#) or ANXA7-GFP overexpression. Arrowheads indicate colocalized TIA1 and DIC1B spots. Scale bars =1 μm (top) and 200 m (bottom). **(I)** Pearson’s coefficient quantifying co-localization between TIA1 and DIC1B from **(H)** (n = 38, 32, 37, 35). **(J)** Western blot analysis detecting Myc-ANXA7 levels in HEK293T cells transfected with shANXA7 (shA7) and either Myc-ANXA7 wild type (Myc-A7) or shRNA-resistant Myc-ANXA7 (Myc-A7-res). 2 μg (large dots) or 1 μg (small dots) of Myc-A7-res plasmid were used for transfection. **(K)** Represented images showing colour-coded GFP-TIA1 lifetime in the soma of transfected neurons, with lifetime **(L)** and FRET efficiency **(M)** quantified and compared across the indicated groups. Scale bars =10 μm (n = 25, 35, 35, 53). Data represent mean ±SEM; in **(I, L, M)** one-way ANOVA. \**p*<0.05, \*\**p*<0.01, \*\*\**p*<0.001, ns non-significant.

**Supplementary Figure 3.**
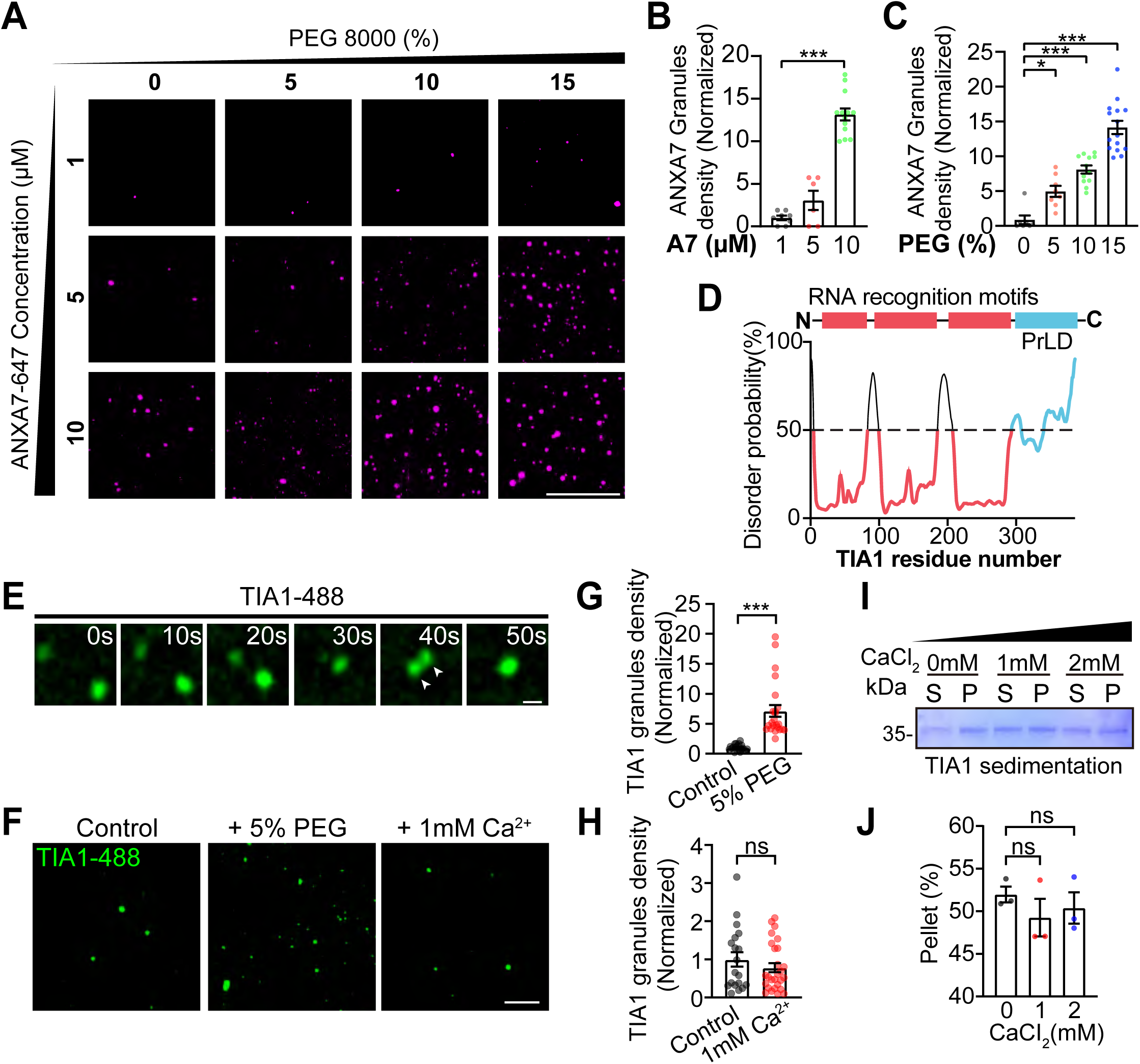
The TIA1 phase separation is not affected by Ca^2+^ elevation. **(A)** *In vitro* assay demonstrating the phase separation of purified ANXA7-647 induced by PEG-8000 addition. The concentrations of PEG and ANXA7-647 are indicated. Scale bar = 50 µm. **(B-C)** Quantification of **(A)**, showing the ANXA7 granule density under different concentrations of protein **(B)** and PEG **(C)** (For **(B)**: PEG concentration = 0%, n = 8, 6, 13; for **(C)**: ANXA7 concentration = 5 µM, n = 7, 8, 12, 15). **(D)** Schematic diagram of the TIA1 protein domain structure with PrDOS analysis, showing the C-terminal PrLD. **(E)** Time-lapse images showing the *in vitro* LLPS process of purified TIA1 protein (TIA1-488), with fusion events of droplets indicated by arrowheads. Scale bar = 2 µm. **(F)** *In vitro* LLPS assay using purified TIA1 protein (TIA1-488), illustrating the effects of adding 5% PEG or 1 mM Ca^2+^ on phase separation. Scale bar = 10 µm. **(G-H)** Quantification of **(F)**, with the effect of 5% PEG **(G)** or 1 mM Ca^2+^ **(H)** on TIA1 granule density compared to those of control groups, respectively (For **(G)**: n = 20, 23; for **(H)**: n = 19, 27). **(I)** *In vitro* sedimentation assay detected by SDS-PAGE showing the distribution of purified TIA1 protein (5μM) in the supernatant (S) and pellet (P) fractions at indicated concentration of CaCl_2_. **(J)** Quantification of the sedimentation assay results in **(I)** (n = 3). Data represent mean ±SEM; in **(G, H)** two-tailed unpaired *t*-test; in **(B, C, J)** one-way ANOVA. \**p*<0.05, \*\*\**p*<0.001, ns non-significant.

**Supplementary Figure 4.**
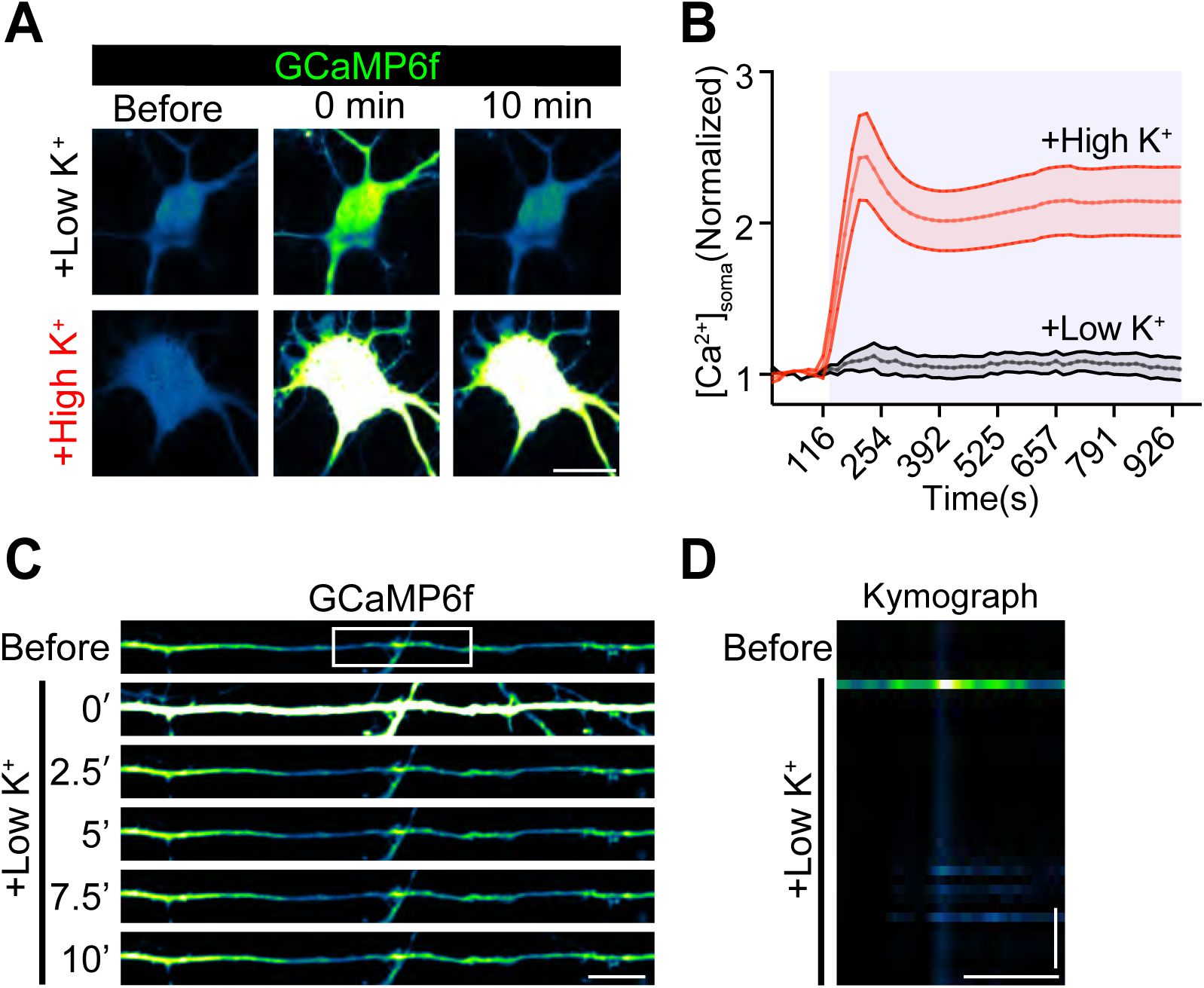
Low K^+^ does not induce Ca^2+^ elevation or ANXA7 aggregation in axons. **(A)** Representative live-images displaying somatic Ca^2+^ concentrations ([Ca^2+^] _soma_) before and after the addition of either 5.6 mM KCl (Low K^+^) or 56 mM KCl (High K^+^) at the indicated time. Scale bar = 20 µm. **(B)** Quantification of the data in **(A)**. Data represent mean ± SEM (“Low K^+^” n = 18, “High K^+^” n = 21). **(C)** Time-lapse images of DIV13 cultured rat hippocampal neurons expressing the Ca^2+^ sensor GCaMP6f. The images show the intensity changes of intracellular Ca^2+^ in the axon following the addition of 5.6 mM KCl (+Low K^+^). Scale bar = 10 µm. **(D)** Kymographs of the axon segment boxed in **(C)**, illustrating the temporal dynamics of Ca^2+^ signaling. Scale Bar = 10 µm; y-axis = 100 s.

**Supplementary Figure 5.**
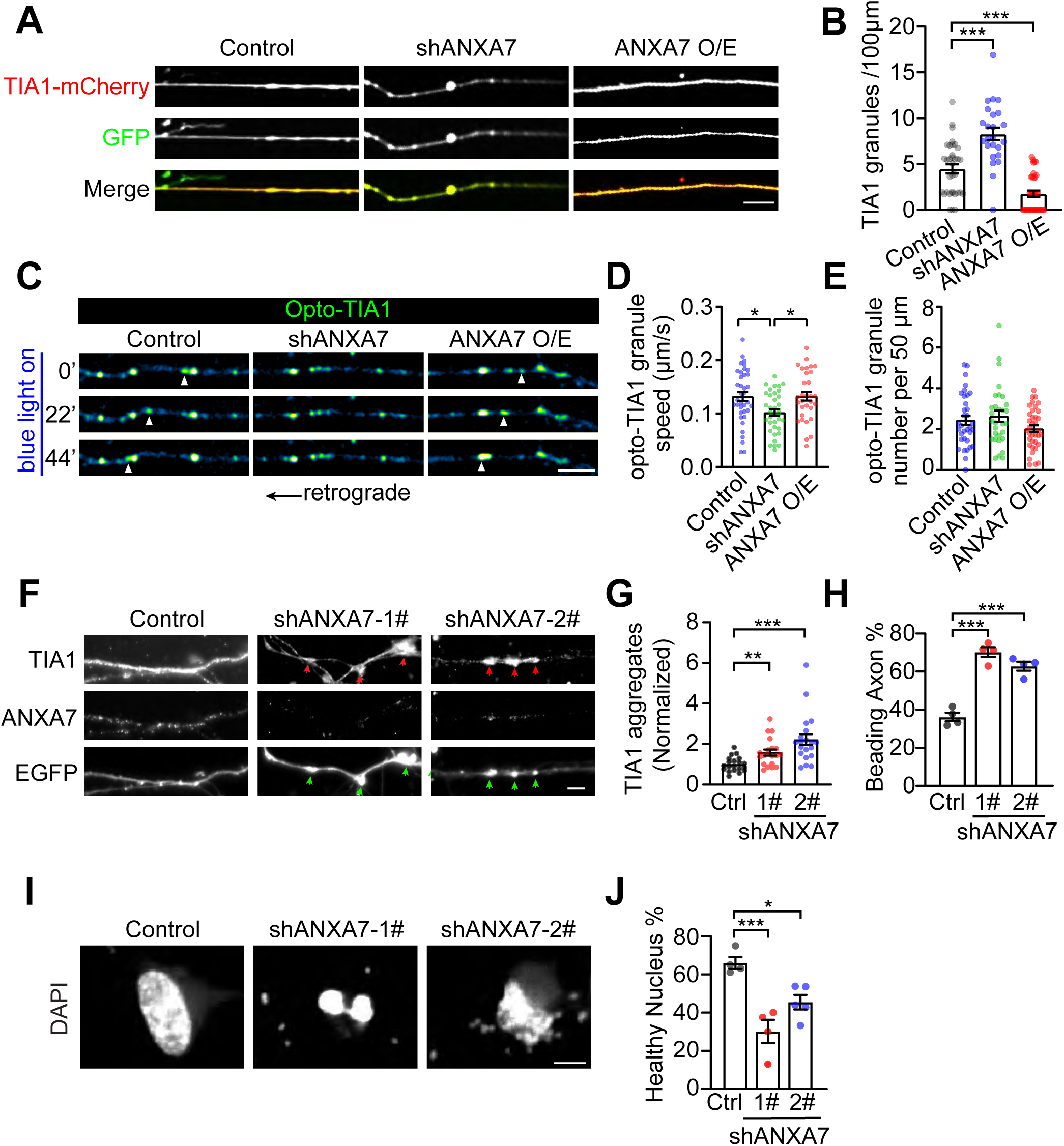
ANXA7 expression regulates TIA1 axon trafficking and phase separation. **(A)** Time-lapse images of TIA1-mCherry in axons of DIV11 rat hippocampal neurons with endogenous ANXA7 knockdown (shANXA7) or ANXA7-GFP overexpression (ANXA7 O/E). Scale bar = 10 µm. **(B)** Quantification of TIA1 aggregates per 100 μm axon under conditions in **(A)** (n = 34, 24, 35). **(C)** Time-lapse images of Opto-TIA1 in DIV9 rat hippocampal neurons with endogenous ANXA7 knocked down (shANXA7) or ANXA7-GFP overexpression (ANXA7 O/E), showing retrograde trafficking of light-induced Opto-TIA1 granules after 120 s blue light exposure. Arrowheads indicate mobile Opto-TIA1 granules, and arrows indicate the retrograde direction. Scale bar = 10 µm. **(D-E)** Quantification from **(C)**, showing the speed **(D)** and density **(E)** of Opto-TIA1 granules in the axons of indicated groups (**(D)**: n = 37, 36, 31; **(E)**: n = 34, 30, 36). **(F)** IF staining images of TIA1 and ANXA7 in axons of DIV9 rat hippocampal neurons with ANXA7 knockdown using two shRNA sequences (shANXA7-1# and 2#). EGFP shows axon morphology, with arrowheads indicating beading structures. Scale bar = 5 µm. **(G-H)** Quantification of **(F)**, showing the density of TIA1 aggregates **(G)** and the ratio of beading axons **(H)** (**(G)**: n = 20; **(H)**: n = 4). **(I)** DAPI staining showing the morphology of neuronal nuclei in DIV9 rat hippocampal neurons with ANXA7 knocked down by shA7-1# and 2#. Scale bar = 5 µm. **(J)** Quantification from **(I)**, showing the proportion of healthy nuclei in the indicated groups (n = 4, 4, 5). Data represent mean ±SEM; in **(G)** two-tailed unpaired *t*-test; in **(B, D, H, J)** one-way ANOVA. \**p*<0.05, \*\**p*<0.01, \*\*\**p*<0.001.

**Supplementary Figure 6.**
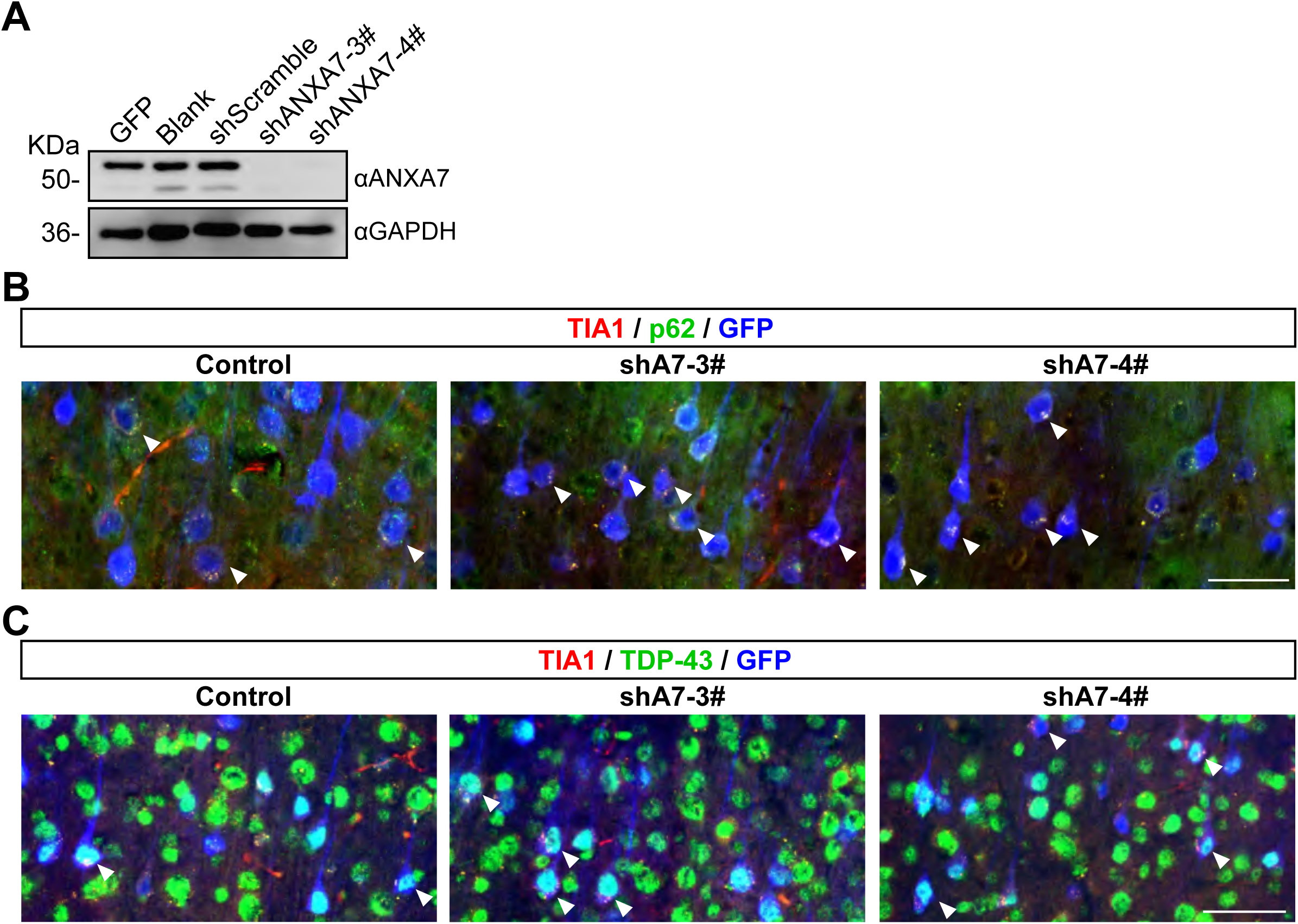
ANXA7 knockdown leads to TIA1 aggregation in layer V neurons of the mouse motor cortex. **(A)** Western blot validating the knockdown efficiency of two different shRNA sequences targeting ANXA7 in mouse brains (shA7-3# and 4#). **(B-C)** Confocal images of endogenous TIA1 and p62 **(B)** or TIA1 and TDP43 **(C)** in layer V neurons in the motor cortex of P60 mice brains infected with AAVs. Infected neurons are identified by GFP expression. Arrowheads indicate neurons containing TIA1 aggregates. Scale bar = 50 µm.

**Supplementary Movie 1. Co-transport of TIA1 granules and RNA along the axon.**

DIV8 hippocampal neurons were transfected with GFP-TIA1 (green) and CY5-UTP (magenta). Time-lapse dual-color confocal microscopy imaging shows the co-transport of TIA1 granules (green) and RNA (magenta), composing the TIA1-containing RNPs along the axons. The region of interest (ROI) is magnified in the right panels. Directions relative to the soma are marked below the magnifications. Scale bar = 50 µm (left), 20 µm (right).

**Supplementary Movie 2. Most TIA1 granules undergo retrograde trafficking in axons.**

DIV8 rat hippocampal neurons cultured in the microfluidic device expressing EGFP-N1 (green) and TIA1-mCherry (magenta) were observed under time-lapse confocal microscopy. Live-imaging shows the directional trafficking of TIA1 granules (magenta) in the axon. The bracketed axonal region is amplified in the bottom panel, with different moving TIA1 granules indicated with arrowheads of different colors. Scale bar = 50 µm (top), 15 µm (bottom).

**Supplementary Movie 3. TIA1 granules are not co-transported with retrograde membranous organelles in axons.**

In DIV8 rat hippocampal neurons cultured in the microfluidic device expressing GFP-TIA1 (green), different retrograde membranous organelles were pulse-chase labelled. Specifically, lysosome (left), endosome (middle), and autophagosomes (right) were labelled using lysotracker, CTB, and BoNT/A-Hc, respectively. Time-lapse confocal microscopy shows the movement of TIA1 granules (green) with each of the retrograde organelles (magenta) in the axon. Scale bar = 5 µm.

**Supplementary Movie 4. TIA1 granules are co-transported with dynein subunit DIC1B.**

In DIV8 rat hippocampal neurons cultured in the microfluidic device expressing GFP-TIA1 and DIC1B-mRFP, time-lapse confocal microscopy shows the co-transport of TIA1 granules (green) and DIC1B (magenta) in the axon. The green hollow arrowheads indicate TIA1 granules, and magenta hollow arrowheads indicate TIA1 co-transported with DIC1B. Scale bar = 5 µm.

**Supplementary Movie 5. Formation of light-induced Opto-TIA1 granules in axons.**

DIV9 rat hippocampal neurons expressing either Opto-Control or Opto-TIA1 were activated with blue light for 120 seconds while time-lapse images were acquired. Representative live images show the formation of Opto-TIA1 granules (bottom) compared to Opto-Control (top). Triangles indicate the light-induced Opto-TIA1 granules. Scale bar = 10 µm.

**Supplementary Movie 6. Light-induced Opto-TIA1 granules undergo rapid retrograde trafficking and fusion in axons.**

DIV9 rat hippocampal neurons expressing Opto-TIA1 were activated with blue light for 65 seconds, with time-lapse images acquired throughout the process. Representative live images show the retrograde trafficking and fusion of Opto-TIA1 granules in the axon. Hollow triangles indicate the movement and fusion of Opto-TIA1 granules. Scale bar = 10 µm.

**Supplementary Movie 7. Opto-TIA1 granules co-transport with ANXA7 granules in axon.**

DIV9 rat hippocampal neurons co-expressing Opto-TIA1 and ANXA7-GFP were activated by blue light while time-lapse images were acquired. Representative time-lapse images show the highly correlated retrograde trafficking of Opto-TIA1 (red) and ANXA7 granules (green) in the axon. Arrows indicate the co-trafficking and fusing processes of Opto-TIA1 and ANXA7-GFP granules. Scale bar = 2 µm.

**Supplementary Movie 8. Time-lapse SIM images showing the co-transported TIA1 and ANXA7 granules in the axon.**

Axon trafficking of DIV9 rat hippocampal neurons co-expressing GFP-TIA1 and ANXA7-mCherry visualized by time-lapse dual-color SIM. The co-trafficking of ANXA7-mCherry (red) and GFP-TIA1 (green) granules in the retrograde direction are indicated with white arrowheads. Scale bar = 5 µm.

**Supplementary Movie 9. High K^+^ depolarization induces Ca^2+^ elevation in neurons.**

In DIV13 neurons expressing GCaMP6f, time-lapse confocal images were acquired to capture the changes in intracellular Ca^2+^ levels induced by the addition of high K^+^ buffer. The yellow background indicates the duration of high K^+^ stimulus, and the bracketed ROI is magnified in the lower panel. Scale bar = 50 µm (top) and 20 µm (bottom).

**Supplementary Movie 10. Ca^2+^ elevation hot spots colocalize with ANXA7 aggregates in axon.**

In axons of DIV13 neurons expressing GCaMP6f and ANXA7-mCherry, dual-colour time-lapse images were acquired to show the formation of Ca^2+^ hot spots (green) and the aggregation of ANXA7-mCherry (red) in response to the addition of high K^+^ buffer. The yellow background indicates the duration of high K^+^ stimulus, and the white arrows represent hot spots. Scale bar = 10 µm.

**Supplementary Movie 11. ANXA7 expression affects the dynamics of TIA1 granules in axons.**

In DIV8 rat hippocampal neurons, GFP-TIA1 was co-transfected with either Myc-ANXA7 (ANXA7 OE) or siANXA7 (ANXA7 KD), and then an axon-segment FRAP assay was conducted to show the dynamics of GFP-TIA1 granules over a long segment of axon shafts. The representative FRAP movie depicts the intensity recovery of GFP-TIA1, demonstrating the molecular dynamics of TIA1 in axons. Scale bar = 10 μm.

**Supplementary Movie 12. ANXA7 expression level affects the trafficking efficiency of TIA1 granules in axons.**

DIV8 rat hippocampal neurons expressing TIA1-mCherry were co-transfected with either ANXA7-GFP (ANXA7 OE) or shANXA7 (ANXA7 KD). Representative time-lapse confocal images depict the axon trafficking of TIA1 granules in the indicated groups. Different TIA1 granules are indicated with different coloured hollow arrows. Scale bar = 5 µm.

**Supplementary Movie 13. Down-regulation of ANXA7 leads to the formation of TIA1 aggregates in axons.**

In DIV8 rat hippocampal neurons, GFP-TIA1 was co-transfected with either control siRNA (siControl) or siANXA7, and then a FRAP assay was conducted on large GFP-TIA1 granules to examine the dynamics of GFP-TIA1 molecules focally. The representative FRAP movie depicts the intensity recovery of GFP-TIA1, demonstrating the molecular dynamics of TIA1 molecules within these large condensates. Scale bar = 2 μm.

**Supplementary Table 1. The interactome of TIA1 and ANXA7 from BioGrid database.**

**Supplementary Table 2. Mass spectrometry analysis of TIA1 and ANXA7 interactome in rat brain lysate.**

**Supplementary Table 3. GO and KEGG analysis of TIA1 and ANXA7 interacting proteins.**

**Supplementary Table 4. RNA sequences used for knocking down ANXA7.**

